# An internally controlled system to study microtubule network diversification links tubulin evolution to the use of distinct microtubule regulators

**DOI:** 10.1101/2024.01.08.573270

**Authors:** Andrew S. Kennard, Katrina B. Velle, Ravi Ranjan, Danae Schulz, Lillian K. Fritz-Laylin

## Abstract

Diverse eukaryotic cells assemble microtubule networks that vary in structure and composition. While we understand how cells build microtubule networks with specialized functions, we do not know how microtubule networks diversify across deep evolutionary timescales. This problem has remained unresolved because most organisms use shared pools of tubulins for multiple networks, making it impossible to trace the evolution of any single network. In contrast, the amoeboflagellate *Naegleria* uses distinct tubulin genes to build distinct microtubule networks: while *Naegleria* builds flagella from conserved tubulins during differentiation, it uses divergent tubulins to build its mitotic spindle. This genetic separation makes for an internally controlled system to study independent microtubule networks in a single organismal and genomic context. To explore the evolution of these microtubule networks, we identified conserved microtubule binding proteins and used transcriptional profiling of mitosis and differentiation to determine which are upregulated during the assembly of each network. Surprisingly, most microtubule binding proteins are upregulated during only one process, suggesting that *Naegleria* uses distinct component pools to specialize its microtubule networks. Furthermore, the divergent residues of mitotic tubulins tend to fall within the binding sites of differentiation-specific microtubule regulators, suggesting that interactions between microtubules and their binding proteins constrain tubulin sequence diversification. We therefore propose a model for cytoskeletal evolution in which pools of microtubule network components constrain and guide the diversification of the entire network, so that the evolution of tubulin is inextricably linked to that of its binding partners.

## INTRODUCTION

The microtubule cytoskeleton is an evolutionarily conserved polymer system that assembles into diverse networks to drive a wide variety of cellular functions. These networks comprise microtubules assembled from α/β-tubulin heterodimers, as well as various microtubule binding proteins. At least two microtubule networks—dynamic mitotic spindles and stable flagellar axonemes—were present in the last common eukaryotic ancestor (Cavalier-Smith, 1978; Mitchell, 2017; Pickett-Heaps, 1974; Pickett-Heaps, 1975), though they have been modified in specific lineages (Bremer et al., 2023; Chaaban & Brouhard, 2017; Heath, 1980; Mitchell, 2017). While the composition and regulation of flagella and mitotic spindles are increasingly well understood in individual species, it is unclear how these and other microtubule networks evolve and diversify across the eukaryotic tree. Because the microtubule cytoskeleton controls so many functions in eukaryotic cells, the diversification of microtubule networks can have major impacts on eukaryotic phenotypes.

In addition to between-species diversity, there is considerable within-species diversity of microtubule networks. Key molecular mechanisms underlying microtubule network diversity within individual cells have been well documented (Akhmanova & Steinmetz, 2015; Goodson & Jonasson, 2018; McNally & Roll-Mecak, 2018; Roll-Mecak, 2020). One such mechanism uses specific subsets of microtubule binding proteins to modulate microtubule network architecture and dynamics. For example, certain families of kinesins are primarily active in the mitotic spindle, including kinesin-5, which promotes spindle bipolarity, and kinesin-7, which facilitates chromosome congression, while other kinesin families are exclusively active in flagella, such as the kinesin-2 family which participates in intraflagellar transport. Non-kinesin microtubule binding proteins can also specialize networks. PRC1, for example, cross-links microtubules in the spindle midzone, whereas a wide class of proteins known as microtubule inner proteins (MIPs) stabilize axonemal microtubules by binding to their inner, lumenal surfaces (Alfieri et al., 2021; Ichikawa et al., 2017; Kajtez et al., 2016; Owa et al., 2019). While microtubule binding proteins represent a powerful mode for achieving specialized microtubule network function, many microtubule binding proteins are “generalists” that function in multiple networks.

Another mechanism used to specialize microtubule networks is to specialize the tubulin subunits themselves. Many eukaryotic genomes encode multiple α- and β-tubulin genes with distinct sequences. The amoeboflagellate *Naegleria gruberi*, for example, uses one set of α- and β-tubulin genes during mitosis, and another set during transient differentiation into a non-dividing flagellate (Chung et al., 2002; Velle et al., 2022). The discovery of multiple tubulin gene loci in *Naegleria* inspired the “multi-tubulin hypothesis,” which posits that distinct tubulins can be used for specific microtubule functions (Fulton & Simpson, 1976). This phenomenon is not limited to *Naegleria:* specific tubulins are also used within the *Drosophila* sperm axoneme, the mitotic spindle of the *Tetrahymena* macronucleus, the unique helical filaments of foraminifera, and neuronal processes in animals (Guo et al., 2011; Hou et al., 2013; Nielsen et al., 2001; Pucciarelli et al., 2012). Tubulins can be further specialized by post-translational modifications—a concept often called the “tubulin code”. These modifications promote differential recruitment of microtubule binding proteins, and may also affect intrinsic microtubule properties (Janke & Magiera, 2020; Roll-Mecak, 2020). Taken together, specialized microtubule binding proteins, the multi-tubulin hypothesis, and the tubulin code provide a foundation for understanding how cells generate distinct microtubule networks within individual cells. However, it is unclear how these intracellular modes of microtubule network specification constrain or guide the diversification of microtubule networks over evolutionary timescales.

Understanding the impact of microtubule binding protein specialization and tubulin diversification on microtubule network evolution is complicated by interconnections between microtubule networks. These interconnections can take many forms. For example, networks can be temporally coupled, as in *Tetrahymena*, where the structures assembled by distinct tubulins coexist in the same cell simultaneously (Pucciarelli et al., 2012). This presents a technical challenge to determine which microtubule binding proteins are specific to which microtubule networks. Networks can also be coupled through shared tubulin pools, as in most animal cells, which assemble distinct networks such as cilia and mitotic spindles from a shared pool of tubulin subunits (Lewis et al., 1987; Lopata & Cleveland, 1987). Thus, the tubulins comprising both cilia and mitotic spindles are subject to evolutionary constraints imposed by both networks (Nielsen et al., 2010). A model system which produces temporally and genetically uncoupled microtubule networks, therefore, may unmask the impact of mechanisms of microtubule network specification on network evolution.

A clear example of a model system with uncoupled microtubule networks is the amoeboflagellate *Naegleria gruberi*. Interphase *Naegleria* amoebae contain no microtubules (Fulton & Dingle, 1971; Velle et al., 2022; Walsh, 2007), but assemble a mitotic spindle *de novo* prior to cell division. *Naegleria* can also exit the cell cycle and transition to a secondary swimming cell type by building flagella and a network of cytoplasmic microtubules specific to the flagellate state. This unique biology separates mitotic and flagellar microtubule networks in both space and time. Furthermore, because distinct tubulin gene loci are expressed during mitosis and differentiation, the pools of tubulin subunits used for these microtubule networks are also genetically uncoupled. Intriguingly, *Naegleria* mitotic tubulins have diverged in sequence relative to other tubulins. While *Naegleria* flagellate tubulins share more than 75% identity with human tubulins, the mitotic tubulins share only 58% identity (Velle et al., 2022). Furthermore, the sequence divergence *between* the mitotic and flagellate tubulins exceeds the variation seen between tubulins of individual animal species: *Naegleria* mitotic and flagellate tubulins share between 57-65% identity while human tubulins all share more than 72% identity. The striking divergence of *Naegleria* mitotic tubulins suggests that intrinsic differences in the tubulin sequences likely contribute to the diversification of *Naegleria* mitotic and flagellar microtubule networks.

The evolutionary distance between *Naegleria* and well-studied model systems also makes it an excellent subject for studying microtubule network evolution. *Naegleria* is part of the eukaryotic supergroup Discoba, which diverged from the Opisthokont lineage of animals and fungi over 1.5 billion years ago (Strassert et al., 2021). These large time scales are sufficient for even highly conserved proteins like tubulins to have diverged, making it possible to examine correlations between unique *Naegleria* biology and cytoskeletal sequence divergence. This approach, however, requires knowing the molecular composition of *Naegleria* microtubule networks, a research area that is largely unexplored. Although a number of genes associated with the flagellate state have been identified by microarray (Fritz-Laylin & Cande, 2010), the sole known component of the mitotic spindle is tubulin. Without knowing the microtubule binding proteins associated with each network, we cannot determine how such proteins specify microtubule network functions, nor how they influence, or are influenced by, mitotic and flagellar tubulin diversity.

Here we explore the roles of microtubule binding proteins and tubulin diversification in specifying the mitotic and flagellar microtubule networks of *Naegleria gruberi*. By comparing microtubule network gene repertoires across a diverse sample of eukaryotes, we confirm that the *Naegleria* genome encodes an extensive complement of microtubule binding proteins. Using transcriptional profiling, we determine that most microtubule binding proteins are upregulated exclusively during mitosis *or* differentiation. By mapping the predicted binding sites of these network components onto *Naegleria* tubulin sequences, we find that residues that interact only with flagellar microtubule network regulators have higher sequence divergence in mitotic tubulins. This correlation suggests that losing a subset of binding interactions facilitates tubulin sequence divergence. Taken together, our analyses suggest that the evolutionary trajectory of tubulins and their binding proteins are interlinked, such that interactions between tubulins and microtubule binding proteins constrain the evolution of the entire network.

## RESULTS

### The *Naegleria* genome encodes a wide variety of conserved microtubule binding proteins

To trace the diversification of microtubule networks across evolutionary timescales, we first need to inventory the microtubule network components found in various eukaryotic lineages. As there is no agreed-upon list of broadly conserved microtubule network components, we developed one using a combination of literature and sequence-based searches (see Methods). We then used this list to identify flagellate and mitotic microtubule network components encoded in the *Naegleria* genome. Microtubule networks typically contain three broad categories of components (**Figure 1, left**): (**1**) *Tubulins and cofactors required for microtubule assembly*. In addition to α- and β-tubulins, this category includes Tubulin Binding Cofactors (Tbc) that assemble tubulin heterodimers (Tian & Cowan, 2013), components of γ-tubulin complexes that nucleate microtubules, and factors that coordinate or activate γ-tubulin localization in the cell, such as augmin, TPX2, and NEDD1 (Kraus et al., 2023). (2) Regulators of microtubule dynamics. This category includes +TIP complex proteins that promote microtubule elongation, the katanin family of AAA ATPases that sever microtubules, as well as microtubule stabilizers, such as the minus-end binding protein CAMSAP/Patronin and microtubule inner-proteins that stabilize axonemal microtubules (Akhmanova & Steinmetz, 2015; Atherton et al., 2019; McNally & Roll-Mecak, 2018; Owa et al., 2019). (3) Components that endow networks with higher-order functions. Microtubule bundling proteins such as PRC1 can generate large-scale organization by cross-linking microtubules, while SSNA1 generates large-scale organization by inducing microtubule branching (Basnet et al., 2018; Kajtez et al., 2016). Additionally, tubulin modifying enzymes generate post-translational modifications leading to differential recruitment of other microtubule binding proteins. Finally, kinesin and dynein motor proteins can promote higher order functions by transporting cargo, crosslinking microtubules, and exerting forces on them. We used well-studied proteins from each of these classes as a foundation for identifying likely microtubule network components in *Naegleria*.

**Figure 1:**
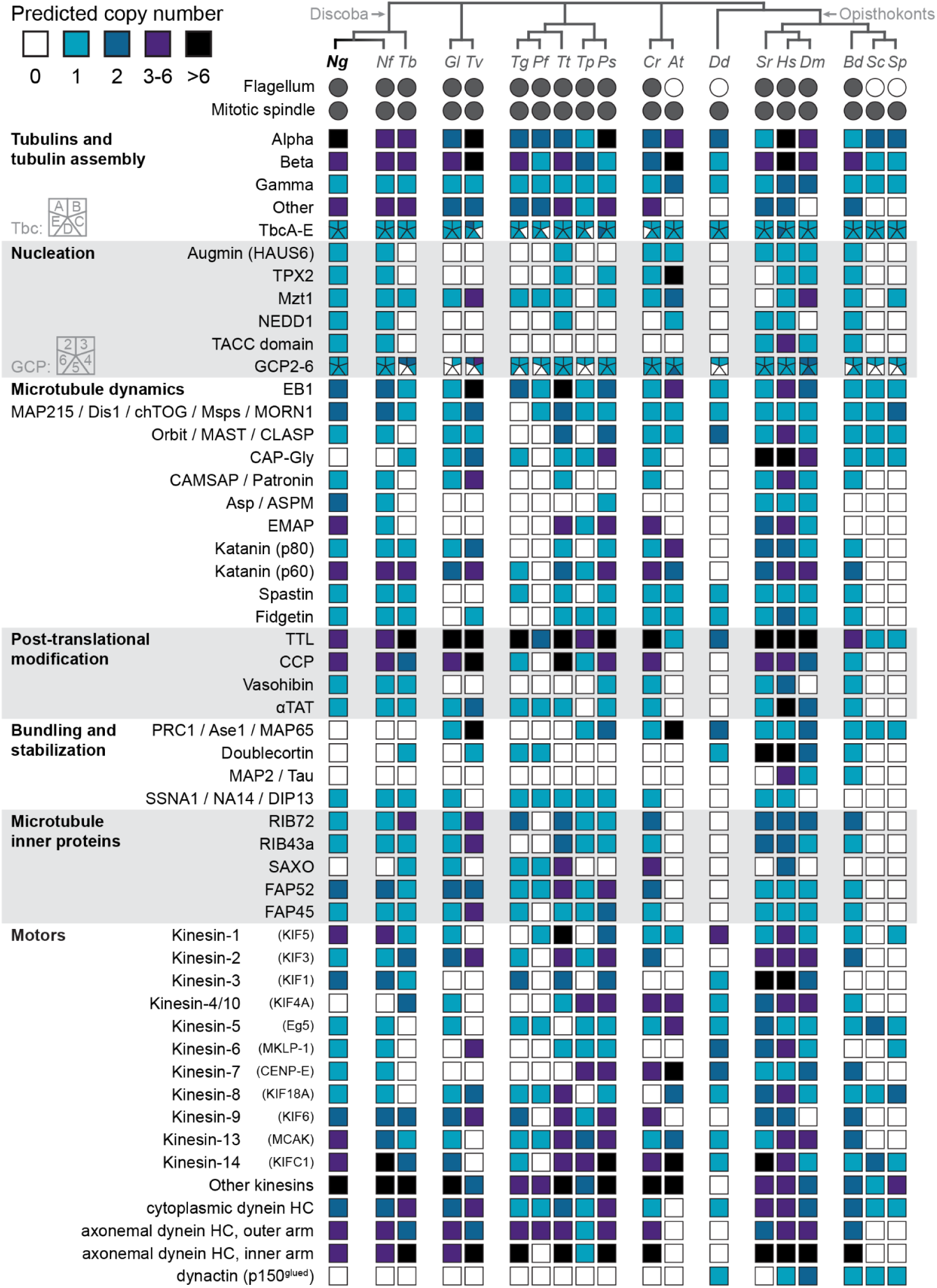
Microtubule network components show patterns of retention and loss across eukaryotes. Distribution of microtubule network components across taxa. Each row represents a different microtubule network component, and each column a different species. Boxes are colored according to the number of homologs of a network component that are predicted in the genome of a given species. Species are organized with a cladogram into major eukaryotic supergroups; from left to right: Discoba; Metamonada; Stramenopiles, Alveolates, and Rhizaria (SAR); Archaeplastida; Amorphea, consisting of Amoebozoa and Opisthokonts. Circles indicate the presence or absence of flagella and mitotic spindles in each species. Species abbreviations, from left to right: *Ng: Naegleria gruberi; Nf: Naegleria fowleri; Tb: Trypanosoma brucei; Gl: Giardia lamblia; Tv: Trichomonas vaginalis; Tg: Toxoplasma gondii; Pf: Plasmodium falciparum; Tt: Tetrahymena thermophila; Tp: Thalassiosira pseudonana; Ps: Phytophthora sojae; Cr: Chlamydomonas reinhardtii; At: Arabidopsis thaliana; Dd: Dictyostelium discoideum; Sr: Salpingoeca rosetta; Hs: Homo sapiens; Dm: Drosophila melanogaster; Bd: Batrachochytrium dendrobatidis; Sc: Saccharomyces cerevisiae; Sp: Schizosaccharomyces pombe. N. gruberi* and *H. sapiens* are bolded to guide the eye. Homologs of 20 network components were only found in one supergroup of eukaryotes and were therefore not displayed, namely: AKAP9, CKAP2, CKAP4/CLIMP-63, GLFND, HURP/DLGAP5, Kar1, MAP1A, MAP1B, MAP6/STOP, MAP7/Ensconsin, MAP8, MAP9, MAP10, MAP70, Msd1/SSX2IP, Mzt2, NuMA, Pericentrin, Spc42, Spc72, and Stathmin. Accession information for the genomes used, and accession numbers and search strategy for identified homologs is listed in **Supplemental Data 1**.

To determine which types of microtubule regulators are broadly conserved, we also identified their homologs in a wide variety of eukaryotic species using a combination of Pfam domain identification, targeted BLAST searches, and phylogenetic analyses (see Methods). Out of 59 classes of microtubule network proteins we investigated, 55 were found in at least four major groups of eukaryotes (**Figure 1**). Of these, 13 classes were nearly universally conserved, including γ-tubulin complex proteins (GCPs), tubulin binding cofactors, and regulators of microtubule dynamics, such as EB1 and MAP215/Dis1 proteins. The remaining 42 classes of microtubule binding proteins were more variably distributed across phyla, with some showing intriguing correlations. For example, we found α-tubulin acetyltransferase (αTAT) only in organisms with flagella, consistent with phylogenetic profiles (Dobbelaere et al., 2023). Furthermore, TPX2 is associated with the augmin subunit HAUS6, consistent with their biochemical interaction in branched microtubule nucleation (Kraus et al., 2023). Taken together, the widespread retention of microtubule regulators across diverse eukaryotic phyla highlights the ancient origin of complex microtubule networks.

Turning to *Naegleria*, we found that its genome encodes 52 of 59 classes of microtubule network components. The missing classes include the dynein cofactor dynactin (which is largely restricted to opisthokonts), the kinesin-4/10 family, which includes chromokinesins that are major contributors to the polar ejection forces in animals that push chromosome arms toward the spindle equator (Almeida & Maiato, 2018), and homologs of the microtubule bundlers and crosslinkers PRC1, Doublecortin, and MAP2/Tau. Additionally, as one might expect, *Naegleria* lacks clear homologs of many well-studied microtubule binding proteins that are only found in plants or metazoa (**Figure 1**, legend). Having identified the potential pool of microtubule regulators in *Naegleria*, we now sought to determine which are used in mitosis and which are used during differentiation.

### Transcriptional profiling of synchronized *Naegleria* differentiation and mitosis

To determine which microtubule network components are used for mitosis and/or differentiation, we began with the tubulins themselves. To confirm previous reports that *Naegleria* transcribes and translates distinct tubulins during differentiation and mitosis (Chung et al., 2002; Velle et al., 2022), we assessed tubulin expression by Western blot using anti-tubulin antibodies. A pan-α-tubulin antibody DM1A labeled bands at ∼54kDa in lysates from dividing amoebae as well as non-dividing flagellates, confirming the presence of tubulin in both populations. To measure the expression of specific tubulins, we used an antibody raised against *Naegleria* flagellar axonemes to target flagellar tubulin (Shea & Walsh, 1987), and a custom antibody raised against mitotic α-tubulin. The flagellate tubulin antibody highlighted a band in lysates from flagellates but not amoebae (**Figure 2A**). Conversely, the custom mitotic tubulin antibody labeled a band in lysates of amoebae but not flagellates (**Figure 2A**). We next used these antibodies for immunofluorescence. While our mitotic tubulin antibody stained only spindles, the flagellate tubulin antibody stained only flagellates (**Figure 2B**). These data support a model in which *Naegleria* microtubule networks are specified, at least in part, by expression of distinct tubulin proteins.

**Figure 2:**
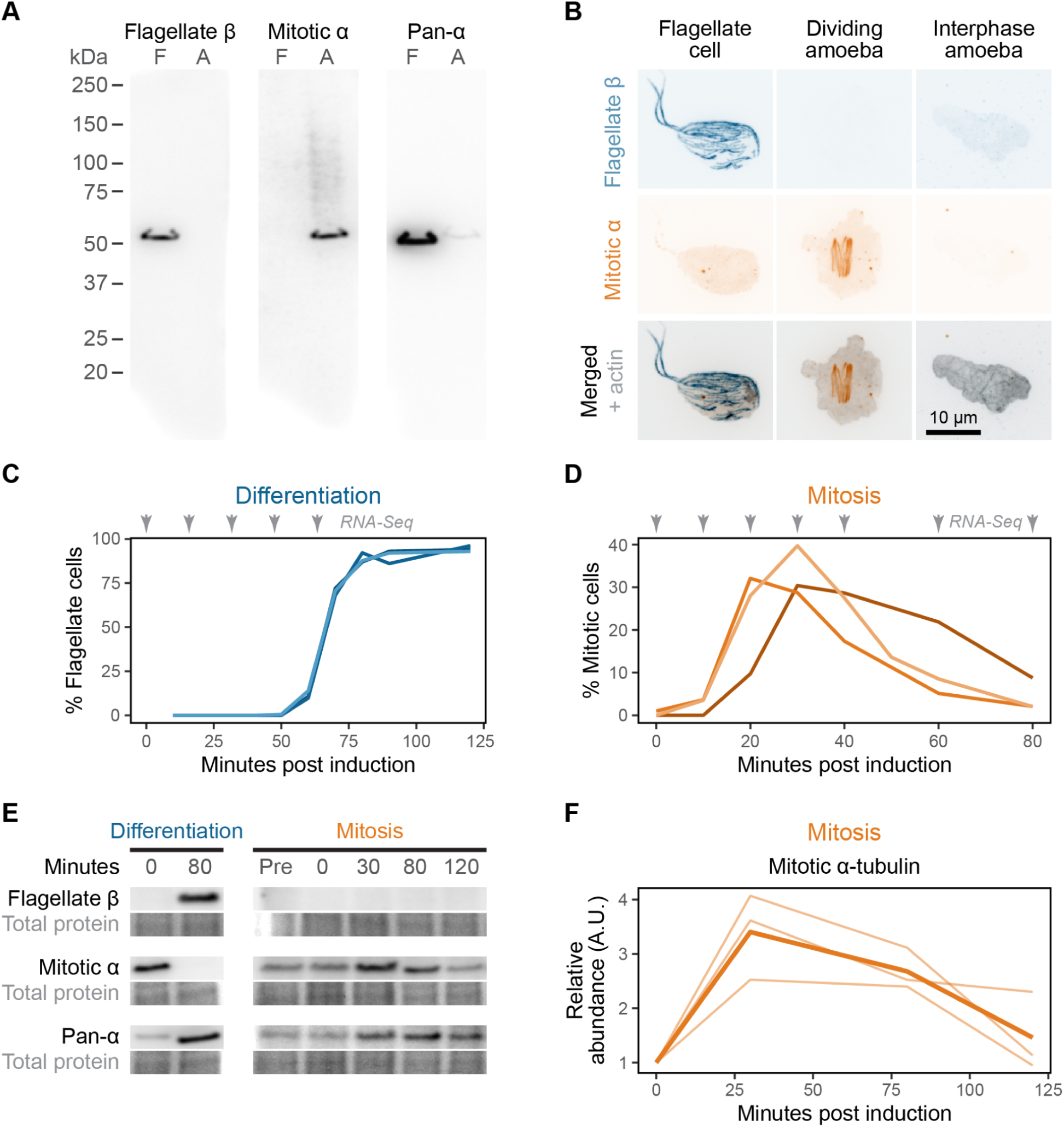
*Naegleria* uses distinct tubulins for synchronized assembly of distinct microtubule networks. **(A)** Western blots of lysates from non-dividing flagellates (marked ‘F’) and dividing amoebae (marked ‘A’), probed with antibodies raised against different tubulin antigens. Flagellate β is a monoclonal antibody raised against *Naegleria* flagellate axonemes; mitotic α is a custom polyclonal raised against a peptide antigen specific to *Naegleria* mitotic α-tubulin; pan-α is the monoclonal α-tubulin antibody DM1A. To enhance contrast, a γ transformation with an exponent of 3 was performed on the image of the pan-α blot. **(B)** Immunofluorescence of cells dual-labeled with the flagellate β and mitotic α antibodies and co-stained with phalloidin to label actin polymer. Scale bar: 10 µm. **(C)** Percentage of flagellates in the population over time after induction of *Naegleria* differentiation. Gray arrows indicate timepoints that were collected for RNA-sequencing. **(D)** Percent of cells with mitotic spindles in the population over time after induction of mitosis. RNA-Seq timepoints are indicated as in (C). **(E)** Western blots of synchronized *Naegleria* differentiation and mitosis, using antibodies as in A. Ponceau S total protein stain is shown as a loading control. **(F)** Quantification of protein abundance during mitotic synchrony shown in (E) for the mitotic α-tubulin antibody. Abundance is shown relative to 0 minutes post induction. Three biological replicate timecourses are shown with thin lines, while the average is shown with a thick line. This figure is connected to **Supplemental Figure S1**.

Having confirmed that *Naegleria* uses different tubulins for mitosis and differentiation, we turned our attention to other microtubule network components. Because *Naegleria* tubulin transcripts are differentially expressed (Chung et al., 2002; Velle et al., 2022), we conducted RNAseq timecourses of synchronized microtubule network assembly to identify which microtubule binding proteins are also upregulated during differentiation and/or mitosis, beginning with differentiation.

The *Naegleria* amoeba-to-flagellate differentiation can be synchronized by washing cells into cold buffer, resulting in over 90% of cells differentiating into flagellates within about an hour (Fulton, 1970). We previously used microarrays to explore transcriptional changes during synchronized differentiation, which revealed upregulation of key microtubule binding protein genes (Fritz-Laylin & Cande, 2010). To extend these analyses to genes not included in the original microarray probe set, and to allow direct comparisons to the mitotic synchrony data detailed below, we processed the same RNA samples for mRNA-Seq with paired-end, short-read sequencing. These samples comprise 3 biological replicates of an 80 minute timecourse sampled every 20 minutes, during which the amoebae fully differentiated into flagellates (**Figure 2C**) (Fritz-Laylin & Cande, 2010). To verify the RNA-Seq data, we compared the relative transcript abundance for each gene as measured by RNA-Seq to our previous microarray analysis of the same samples. Overall, there was strong global correlation between abundances measured by both approaches (**Supplemental Figure S1A**).

We next synchronized the assembly of *Naegleria* mitotic spindles by optimizing the use of heat-shock to arrest the *Naegleria* cell cycle (Fulton & Guerrini, 1969). To track cell cycle progression in real time, we measured cell size and concentration, and detected a synchronized pulse of cell division between 30 and 80 minutes after release from heat shock. By measuring the proportion of cells with mitotic spindles using immunofluorescence (*i.e.* the mitotic index), we observed a 3-fold enrichment in mitotic cells compared to non-synchronized populations, with ∼30% of cells undergoing mitosis at any one time (**Figure 2D**). We confirmed with Western blots that mitotic tubulin protein was enriched during the peak of mitotic synchrony and returned to baseline abundance two hours after release from heat shock. (**Figure 2E, F, Supplemental Figure S1B**). To measure gene expression during mitosis, we collected and sequenced RNA from five biological replicates comprising nine timepoints across the mitotic synchrony experiment: prior to heat shock, halfway through the heat shock, and at seven timepoints between 0 and 80 minutes after release from heat shock.

To assess the biological reproducibility of synchronized differentiation and mitosis we conducted principal component analyses of normalized gene expression and observed tight clustering of biological replicates (**Figure 3A**). For the mitosis dataset, we noted that the timepoints form a contiguous and cyclical trajectory through the space formed by the first two PCA axes (**Figure 3A**). This cyclical trajectory is consistent with the periodic nature of mitosis. Moreover the contiguity of timepoints within both PCA plots suggests that we sampled at a sufficiently high frequency to capture relevant gene expression dynamics during both experiments.

**Figure 3:**
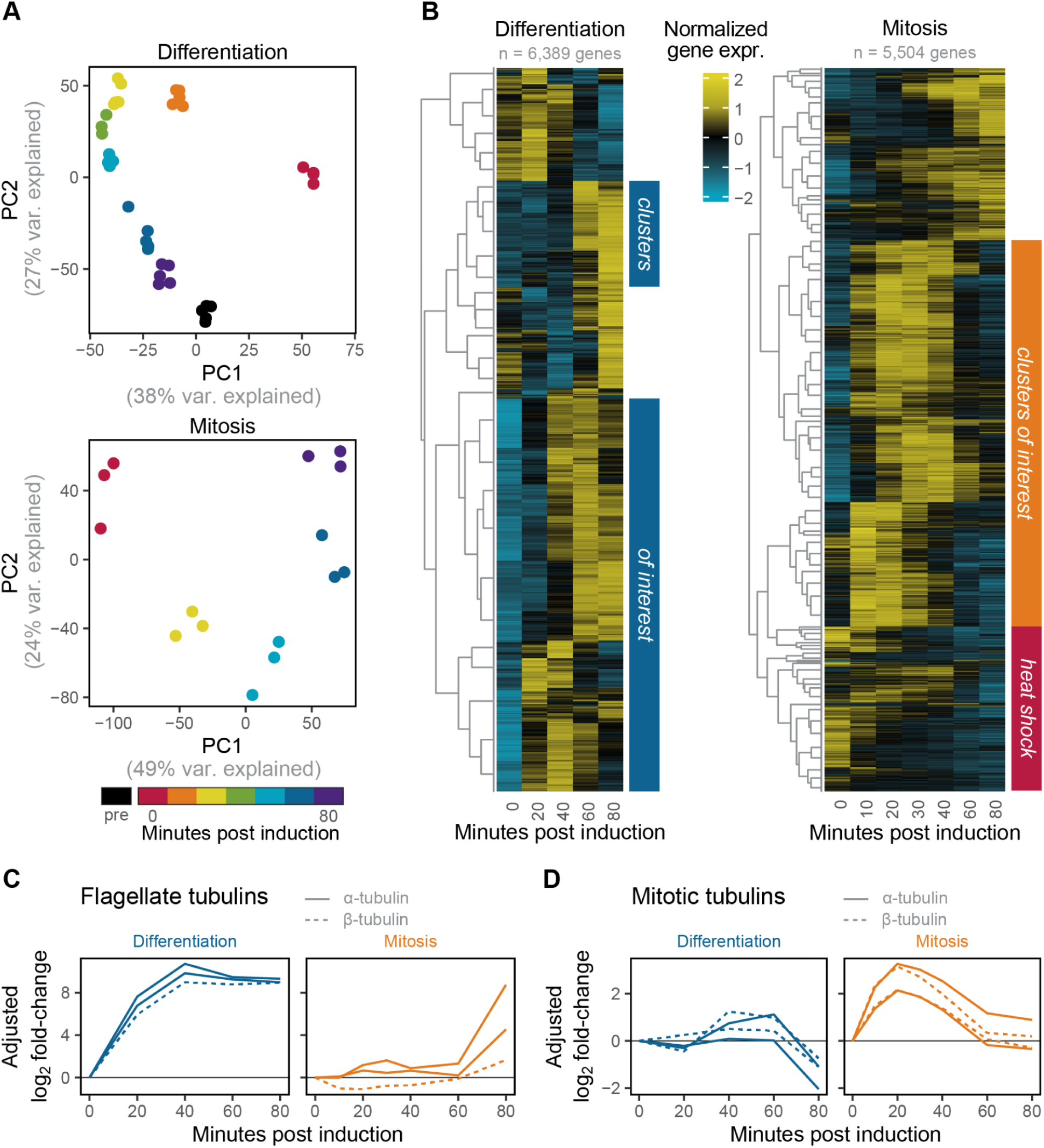
Transcriptional profiling of *Naegleria* differentiation and mitosis. **(A)** Principal component analysis (PCA) of RNA-Seq biological replicates from synchronized differentiation (top) and mitosis (bottom). Colors indicate minutes post induction of differentiation or mitosis. The variance explained along each principal component is shown in gray for each axis. **(B)** Heatmaps of differentially expressed genes in each experiment, clustered into groups. Each row is a gene, and each column is a different timepoint in the experiment. Expression levels are normalized per-gene to have a mean expression level of 0 and standard deviation of 1. Genes clusters based on expression patterns are grouped by hierarchical clustering; gray cladograms on the left of each heatmap indicate the grouping of each cluster, so each branch points to a cluster of multiple genes. Blue colored bars represent clusters we define as differentiation-specific based on their expression profile; orange bars represent mitotic-specific genes. Expression data is provided in **Supplemental Data 2**. **(C-D)** Expression of flagellate tubulins (C) and mitotic tubulins (D) during differentiation and mitotic synchrony. Each line indicates the adjusted log_2_ fold-change of a single gene over time, relative to 0 minutes post induction (see Methods for description of fold-change adjustment). Blue lines show the expression of these genes during differentiation; orange lines show the expression of the same genes during mitotic synchrony. Solid lines are α-tubulins, dashed lines are β-tubulins. Note that the transcript abundance of flagellate tubulins in the mitotic synchrony experiment is near the lower limit of detection; thus the large fold-change increase for these genes at 80 minutes post induction of mitosis still reflects a very small absolute transcript abundance. This figure is connected to **Supplemental Figure S2**.

### Most *Naegleria* microtubule network components are expressed exclusively during differentiation or mitosis

To identify genes used in flagellar and/or mitotic microtubule networks, we next identified which genes are upregulated in each RNA-seq dataset. We found 6,389 genes in the differentiation experiment and 5,504 genes in the mitosis experiment that were differentially expressed more than 2-fold relative to the initial timepoint and that passed the threshold for statistical significance (see Methods). We then clustered these differentially expressed genes using a time-series-aware clustering algorithm, resulting in 29 differentiation clusters and 81 mitosis clusters (**Figure 3B, Supplemental Figure S2A,C,D**). To identify clusters containing microtubule network components, we used the expression of mitotic and flagellate tubulins as a guide. The flagellate tubulins were rapidly induced during differentiation but were not differentially expressed during mitosis (**Figure 3C**). Conversely, mitotic tubulins were not upregulated during differentiation but were maximally upregulated 30 minutes post induction of mitosis, with expression strongly decreasing by 80 minutes (**Figure 3D**). To ensure that our mitotic clusters did not include genes associated with the heat shock response, we investigated the expression of heat shock proteins (**Supplemental Figure S2B**). These genes were strongly upregulated during heat shock, but rapidly *decreased* in expression after release from heat shock, distinct from the mitotic tubulins (**Supplemental Figure S2D**). We therefore defined “differentiation” gene clusters as those with maximal expression after the initial timepoint and “mitotic” gene clusters as those with maximal expression 10-60 minutes after induction (see Methods). Using these definitions, most of the microtubule network components we previously identified (**Figure 1**) are expressed during assembly of flagellar and/or mitotic microtubule networks (**Figure 4A**).

**Figure 4:**
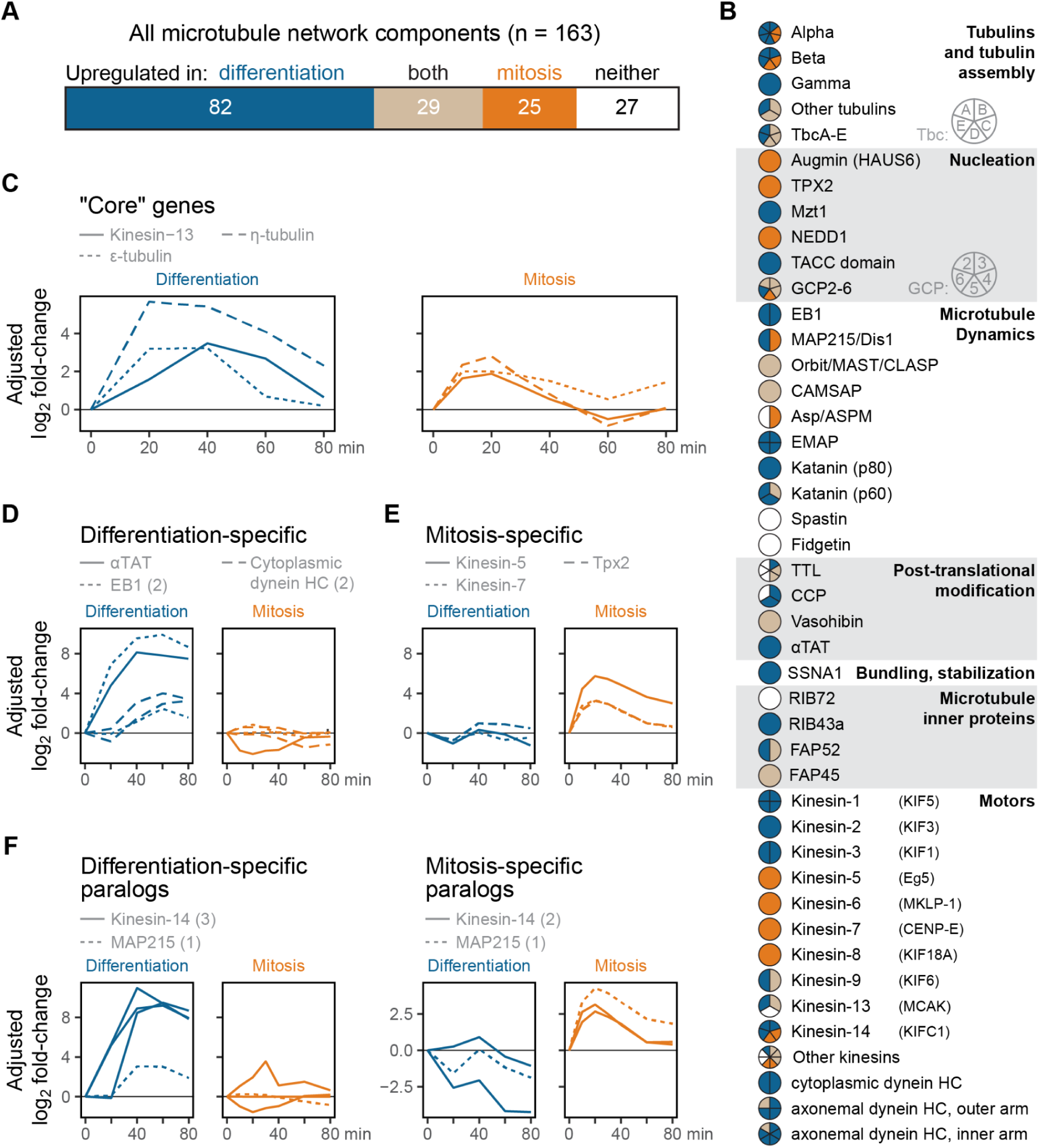
*Naegleria* microtubule network components are largely process-specific. **(A)** Categorization of *Naegleria* microtubule network components as upregulated in differentiation, mitosis, both, or neither. Numbers indicate the number of genes in each category, and bars are proportional to each number. **(B)** Expression of each family of microtubule network components, as in Figure 1. Circles are divided into sectors for each homolog for multi-copy gene families. Color indicates if that homolog is upregulated in differentiation (blue) or mitosis (orange). Only components with predicted *Naegleria* homologs are shown. Accession numbers of each homolog are included in **Supplemental Data 1**, and the categorizations of each gene are provided in **Supplemental Data 2**. **(C-F)** Expression over time of genes significantly upregulated in both differentiation and mitosis (C), only differentiation (D), or only mitosis (E). The genes in (F) are part of multi-gene families in which one set of paralogs is expressed only in differentiation (left) and the other set only in mitosis (right). The adjusted log_2_ fold-change over time for each of these genes is shown, relative to 0 minutes post induction. Blue lines show expression of these genes during differentiation, and orange lines show the expression of the same genes during mitotic synchrony. The line style shows the indicated gene(s), and numbers in parentheses the number of genes in that family that are displayed. See Methods for description of fold-change adjustment. This figure is connected to **Supplemental Figure S3**.

When considering genes that are upregulated in differentiation and/or mitosis, four expression categories naturally emerge: “core” genes upregulated in both differentiation *and* mitosis, “process-specific” genes upregulated in either differentiation *or* mitosis, and genes which are not upregulated in either process. We therefore classified *Naegleria* microtubule binding protein genes into these categories (**Figure 4B**).

Surprisingly, the core microtubule binding proteins include members of the tubulin-binding cofactor (Tbc) complex, which assembles α/β-tubulin heterodimers (**Figure 4B**). The upregulation of Tbc components in both processes suggests that, despite the extensive sequence divergence of *Naegleria* flagellate and mitotic tubulins, they still share a single Tbc complex. Intriguingly, Naegleria homologs of ε- and η-tubulin were upregulated in both differentiation and mitosis. While these tubulin families have been associated with centriole assembly in *Chlamydomonas*, *Paramecium*, and animals (Chang et al., 2003; Dutcher et al., 2002; Ruiz et al., 2000), their upregulation in mitosis may suggest a role for these tubulins in acentrosomal microtubule nucleation in *Naegleria*, as has been found in *Xenopus* extracts (Chang et al., 2003). The microtubule-depolymerizing kinesin-13, which regulates the lengths of spindle and flagellar microtubules of other species (Dawson et al., 2007), is also upregulated during assembly of both microtubule arrays (**Figure 4C**).

Differentiation-specific genes include genes known to be essential for basal-body and flagellar structure, including 5 of 6 inner-arm dyneins, 3 of 4 outer-arm dyneins, and basal body components (e.g. PACRG1, Sas-4, and Sas-6) (**Supplemental Figure S3A**). Surprisingly, several components of the axonemal central pair complex, including the scaffolding proteins CPC1/SPEF2 and PF16/Spag6, are upregulated in mitosis as well as differentiation (**Supplemental Figure S3A**). Also upregulated only in differentiation is the microtubule acetyltransferase αTAT, whose activity is associated with stable microtubules (**Figure 4D**). Accordingly, we previously detected acetylated microtubules in *Naegleria* flagellate cells, but not in mitotic spindles (Velle et al., 2022). This category of genes also includes EB1 and cytoplasmic dynein (**Figure 4D**), both of which play key roles in interphase *and* mitotic microtubule networks in other species (Dema et al., 2022; Gassmann, 2023).

Mitosis-specific genes (**Figure 4E**) include key factors that regulate mitotic spindle assembly in other systems, including members of the kinesin-5 family, which promotes spindle assembly in many animals, fungi, and plants (Bannigan et al., 2007; Ferenz et al., 2010), and the kinesin-7 family, which promotes chromosome congression and kinetochore attachment in animals (Craske & Welburn, 2020). The mitosis-specific genes also include a *Naegleria* TPX2 homolog, consistent with the roles of TPX2 in nucleation of branched microtubules in mitotic spindles of vertebrate cells (Petry et al., 2013).

Finally, given that *Naegleria* tubulins are transcriptionally restricted to differentiation *or* mitosis, we wondered if any other classes of microtubule network components also displayed the same pattern of paralog specialization. By examining the expression patterns of multi-copy microtubule binding proteins, we found two additional examples of this trend: the kinesin-14 family of minus-end directed microtubule motors, and the MAP215/Dis1 family of TOG-domain-containing microtubule polymerases. *Naegleria* has five kinesin-14 paralogs (**Figure 1**), two of which are upregulated exclusively in mitosis while the other three are upregulated only during differentiation (**Figure 4F**). While four of these kinesin-14 paralogs have positively charged N-terminal tails that may facilitates cross-linking to the negatively charged microtubule surfaces (Lüdecke et al., 2018), one *Naegleria* mitotic kinesin-14 has a negatively-charged tail domain (**Supplemental Figure S3B**); this may reflect an impaired ability to cross-link microtubules, or may be related to a reduced negative charge density in mitotic tubulins (Velle et al., 2022). *Naegleria* also has two MAP215 homologs, one expressed in mitosis and the other during differentiation (**Figure 4F**). Like other MAP215 homologs, these proteins contain a tandem array of TOG domains that bind tubulin (**Supplemental Figure S3C,D**) (Ayaz et al., 2012). Interestingly, key residues that are required for binding to yeast tubulin, and that are generally highly conserved, are divergent in the mitotic MAP215 paralog (**Supplemental Figure S3E**), perhaps to maintain interactions with divergent residues in the mitotic tubulins. Overall, this suggests that in addition to tubulins, kinesin-14 and MAP215 families may have undergone duplication and then adopted mitosis-specific and differentiation-specific functions, with increased divergence in mitotic paralogs.

Taken together, these data show that most *Naegleria* microtubule-associated proteins are expressed during either mitosis *or* differentiation, suggesting that they are associated with a single microtubule network. This finding is consistent with a model in which *Naegleria* microtubule network specification relies on differential use of microtubule binding proteins, along with tubulin specialization.

### Mitotic tubulin sequence divergence correlates with loss of binding to differentiation-specific regulators

Because most microtubule network components were upregulated only during differentiation *or* during mitosis, we wondered whether the use of distinct repertoires of microtubule binding proteins in the two networks may be related to the divergence of the mitotic tubulins. This relationship could manifest in different ways. For example, mitotic tubulin could be more divergent at sites that contact mitosis-specific microtubule binding proteins. This possibility would hint at coevolution between mitotic tubulins and mitosis-specific regulators. Alternatively, mitotic tubulin could be more divergent at sites bound by flagellate-specific regulators that are not contacted in the mitotic microtubule network. This second possibility would suggest that interactions with microtubule binding proteins constrain tubulin diversification.

To distinguish between these possibilities, we compared mitotic tubulin divergence at the predicted binding interfaces of differentiation-specific and mitosis-specific microtubule network components. To measure tubulin sequence divergence, we used the TwinCons metric (Penev et al., 2021), which quantifies amino acid similarity between two specified sequence subsets for each position of a multiple sequence alignment. Large positive TwinCons scores reflect conservation in both sequence subsets, negative scores reflect conservation within subsets but divergence between subsets, and scores near zero indicate low conservation in one or both of the sequence subsets (**Figure 5A**). We calculated TwinCons scores based on a multiple sequence alignment of tubulins from five Heterolobosean species—the lineage that includes *Naegleria* and related amoeboflagellates (**Supplemental Figure S4A,B**). We then categorized the sequences as “mitotic” or “flagellate” by phylogenetic analysis (see Methods) and used these categories and the multiple sequence alignment to calculate TwinCons scores. The resulting score distribution across both α- and β-tubulins was bimodal, comprising a tight distribution of highly conserved residues, and a broad distribution of more weakly conserved residues (**Figure 5B**). Inspection of the alignment suggested that the highly conserved residues were conserved in both flagellate and mitotic tubulins, while weakly conserved residues tended to be conserved within flagellate tubulins but not within mitotic tubulins (**Supplemental Figure S4A,B**). These data are consistent with previous findings that the mitotic tubulins have diverged relative to flagellate tubulins as well as tubulins of other eukaryotic species (Chung et al., 2002; Velle et al., 2022). We then identified tubulin residues that interact with specific microtubule binding proteins. We examined 29 molecular structures of microtubule binding proteins, which were solved in complex with microtubules, and which have a *Naegleria* homolog (**Supplemental Data 3**). To determine which tubulin residues interface with a given microtubule binding protein, we identified the amino acid residues of tubulin that have a solvent-exposed area that is predicted to decrease when bound to that protein (see Methods).

**Figure 5:**
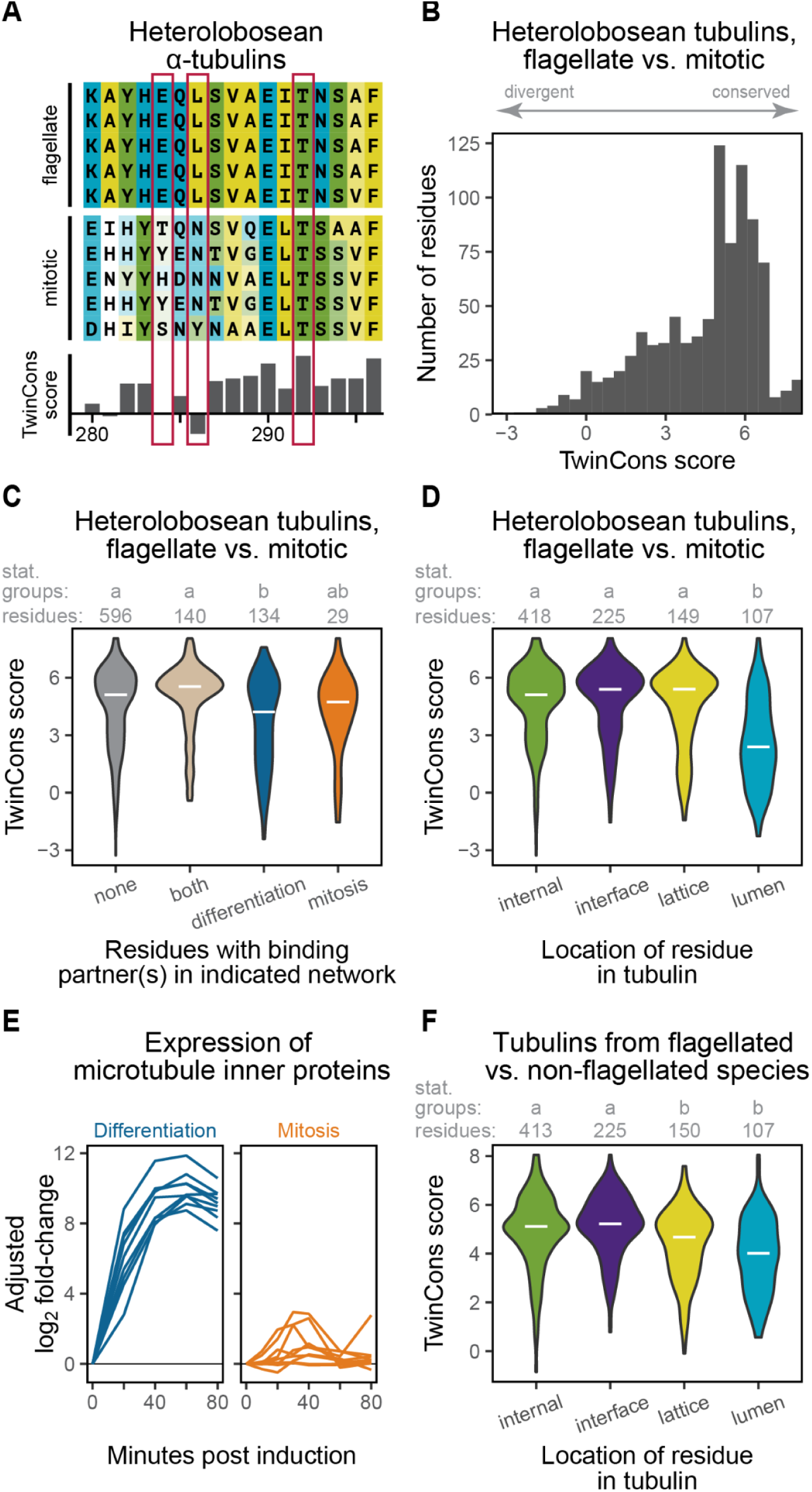
Divergent residues in *Naegleria* mitotic tubulins co-occur with differentiation-specific regulator binding sites. **(A)** Example alignment of α-tubulins from Heterolobosean species with the calculated TwinCons score below. Residues are colored based on their relative hydrophobicity, and numbered based on the homologous positions in pig α-tubulin. Examples (from left to right) of alignment positions with TwinCons scores near zero, negative, and strongly positive are highlighted with red boxes. **(B)** Distribution of TwinCons scores across tubulin residues. TwinCons scores measure per-residue conservation compared between two sequence subsets of a multiple sequence alignment, in this case mitotic vs. flagellate tubulins of *Naegleria* and its relatives. Positive scores indicate residues that are conserved across both subsets of the alignment, while scores near zero indicate weak conservation in at least one subset of the alignment. Negative scores indicate residues that are conserved within subsets but divergent between subsets. The alignments of tubulins used to calculate the TwinCons scores are provided in **Supplemental Figure S4A-B**. **(C)** Distributions of TwinCons scores as in (B), partitioned based on whether residues are predicted to bind to the indicated category of microtubule regulators. Shared residues bind to both differentiation- and mitotic-specific regulators, while “mitosis only” and “differentiation only” residues only bind to regulators specific to those processes. The number of residues in each group is listed below each violin plot. White lines indicate the median of the distribution. Letters indicate statistically distinguishable distributions. **(D)** Distribution of TwinCons scores as in A, partitioned based on residue location within the microtubule: internal to the tubulin monomer, at the lateral or longitudinal interfaces between microtubule protofilaments, on the microtubules’ external surface (lattice), or on the internal surface (lumen). Letters indicate statistically distinguishable distributions (STATS). **(E)** Adjusted log_2_ fold-changes in gene expression (see Methods) for microtubule inner proteins over time. Blue lines show expression during differentiation, and orange lines show expression during mitosis. **(F)** TwinCons scores comparing tubulins from flagellated species to those from non-flagellated species (*Naegleria* is excluded from this analysis). The scores are partitioned based on residue location in the microtubule as in (D). Letters indicate statistically distinguishable distributions . The alignments used to calculate these TwinCons scores are provided in **Supplemental Figure S4C-D**. Statistically distinguishable groups in all panels are defined by Dunn’s post-hoc Bonferroni-corrected test, with p < 0.001. This figure is connected to **Supplemental Figure S4**.

Having calculated the conservation of each tubulin residue and predicted which residues bind to other microtubule network components, we next correlated these two features. Because some tubulin residues bind multiple microtubule binding proteins, we partitioned tubulin residues based on whether they bind: only differentiation-specific microtubule network components, only mitosis-specific network components, or both. We found a significant difference in TwinCons scores between all three groups (Kruskal-Wallis test, chi-squared = 26.1007, df = 3, p < 1e-5), with “differentiation-only” residues tending to be less conserved (Dunn’s post-hoc Bonferroni-corrected test, p < 0.001) (**Figure 5C**). This divergence of “differentiation-only” residues suggests that mitotic tubulins may have diverged after gene duplication and specialization because they were released from constraints imposed by differentiation-specific microtubule binding proteins.

The number and diversity of microtubule binding proteins whose structure has been solved in complex with microtubules, however, is limited. We therefore complemented our structure-based method with a more holistic approach by classifying tubulin amino acid residues based their location in the microtubule: on the outer surface (lattice), inner surface (lumen), at the interface between tubulin subunits, and internal to the monomer structure. We found that lumenal residues were more divergent than residues in other parts of the microtubule (Kruskal-Wallis test, chi-squared = 79.237, df = 3, p < 1e-5, with Dunn’s post-hoc Bonferroni-corrected test p < 1e-5) (**Figure 5D**). Intriguingly, cryo-electron tomography has identified a growing number of microtubule inner proteins (MIPs) that bind lumenal residues and stabilize microtubules of the axoneme (Ichikawa et al., 2017; Owa et al., 2019). We therefore wondered whether divergence of lumenal residues may be driven by differential use of lumen-binding proteins in each microtubule network. Accordingly, we explored the expression of *Naegleria* MIP homologs and found that they are all exclusively upregulated during differentiation (**Figure 5E**). Taken together, these data suggest that increased divergence of lumenal mitotic tubulin residues may be due to a loss of constraint imposed by differentiation-specific MIPs.

If the lumenal divergence we observe in *Naegleria* mitotic tubulins is due to loss of MIP-imposed constraints, then we might expect to find similar divergence in tubulins from other species that no longer build flagellar microtubule networks. Indeed, previous analyses identified specific tubulin motifs involved in binding to MIPs that are often absent in organisms that lack flagella (Ichikawa et al., 2017). To directly test this hypothesis, we therefore used TwinCons analysis to compare tubulins from organisms that have flagella to those that lack flagella. We again found that lumenal residues were significantly less conserved than residues from other microtubule locations (Kruskal-Wallis test, chi-squared = 61.947, df = 3, p < 1e-5; Dunn’s post-hoc test p < 0.001) (**Figure 5F**). Inspection of the alignment confirmed that this effect was driven by increased divergence in sequences from species lacking flagella (**Supplemental Figure S4C,D**), supporting the idea that loss of flagellate microtubule network components is associated with increased tubulin sequence divergence. Taken together, these data suggest that network-specific repertoires of microtubule binding proteins constrain and guide tubulin sequence evolution.

## DISCUSSION

Here we show that *Naegleria* expresses two different pools of microtubule network components, one during assembly of mitotic spindles and the other during flagellate differentiation. Each pool comprises its own distinct tubulins and microtubule binding proteins. We also uncover a correlation between sites of *Naegleria* tubulin divergence and sites that bind to proteins found in a single pool of regulators. From this correlation we infer that the use of distinct tubulins and different microtubule binding proteins both contribute to functional specification of microtubule networks in *Naegleria*.

Our analyses also include full genome transcriptional profiling of flagellate and mitotic microtubule network assembly. *Naegleria* was the first documented case of *de novo* basal body assembly, and these data confirm and extend prior work showing that *Naegleria* expresses basal body and axonemal proteins just prior to formation of its flagellar microtubule array (Fritz-Laylin & Cande, 2010). We also identify 54 microtubule network components associated with mitosis. This is a major step toward understanding *Naegleria* cell division, as tubulin was the only previously known spindle component in this system (Velle et al., 2022). These mitotic network components include kinesin-5 and kinesin-14, which promote spindle bipolarity in animals and yeast through antagonistic activity (Mountain et al., 1999; Pidoux et al., 1996).

Their upregulation during *Naegleria* mitosis suggests that such a mechanism may also be present here. Intriguingly, TPX2 and the augmin subunit HAUS6 are also upregulated during mitosis, suggesting that branched microtubule nucleation may be a source of microtubules in the *Naegleria* spindle, as for other systems. Branched nucleation could explain how *Naegleria* assembles pronounced microtubule bundles in the absence of centrioles or other obvious microtubule organizing centers (Velle et al., 2022). Although we identify a large number of mitotic regulators shared with other systems (**Figure 4B,C**), we did not detect upregulation of some known mitotic regulators like EB1 or cytoplasmic dynein (Dema et al., 2022; Gassmann, 2023). This may reflect biological differences for these regulators in *Naegleria* mitosis.

Alternatively, their constitutive expression may be sufficient to support mitotic function, or their upregulation may be obscured by incomplete synchrony in our experiment. We also found several structural components of the central pair complex that were unexpectedly upregulated in both mitosis and differentiation (**Supplemental Figure S3A**). Their mitotic expression may hint at unique microtubule organization in *Naegleria* mitosis. Although this could also be explained by a small subpopulation of differentiating cells, we find this explanation unlikely because most other axonemal components are not expressed during mitosis. In any event, among the microtubule network components that *are* upregulated in mitosis, we find two gene families that—like mitotic tubulins—have at least one paralog that appears specific to mitosis. These mitosis-specific paralogs have divergent properties relative to those of other species as well as the differentiation-specific paralogs (**Supplemental Figure S3**). These mitotic-specific paralogs, therefore, may represent examples where microtubule regulators coevolve with network-specific tubulins. Overall, the distinct transcriptional programs associated with the assembly of *Naegleria* flagellate and mitotic microtubule networks highlights the potential for network diversification by incorporation of specialized network components.

We also confirm striking differences between *Naegleria* flagellate and mitotic tubulin sequences. The substantial divergence of *Naegleria* mitotic tubulins may alter their susceptibility to commonly-used microtubule inhibitors. We previously found that *Naegleria* mitosis is insensitive to a wide range of small-molecule inhibitors (Velle et al., 2022), including the microtubule stabilizing drug taxol. Moreover, we found that taxol can bind to *Naegleria* flagellate microtubules but not mitotic microtubules (Velle et al., 2022). The proximity between the binding sites of taxol and microtubule inner proteins—both of which bind within the microtubule lumen—suggests that the disruption of one could entail the loss of the other. Accordingly, other organisms that have lost flagella appear insensitive to taxol, including *Saccharomyces* (Gupta et al., 2003). We therefore hypothesize that constraints imposed by specific microtubule binding partners may predict the availability of pharmacological targets for microtubule disruption. Such sequence divergence within tubulins could be advantageous for escaping the microtubule-disrupting activities of antagonistic species (Forli, 2014; Higashide et al., 1977; Lee et al., 2012).

In addition to identifying microtubule network components in *Naegleria*, we examined their conservation across major eukaryotic phyla. This analysis builds upon previous studies that either traced the conservation of specific types of microtubule network components (Dacheux et al., 2012; Gard et al., 2004; Wickstead et al., 2010; Wickstead & Gull, 2007), or examined the conservation of many network components within a single eukaryotic group (Bodakuntla et al., 2019; Gardiner, 2013). Here we examined the conservation of microtubule binding protein families across eukaryotic lineages using a single, consistent strategy for homolog identification. Although our conservative approach readily identified dozens of microtubule binding proteins in many species, it is possible that some homologs may have escaped detection. Moreover, the list of known microtubule network components is almost certainly missing proteins that are widely conserved, but not found in humans and other heavily-studied species. Despite these limitations, the conservation of so many microtubule regulators in species that diverged at the dawn of eukaryotic evolution underscores the complexity of ancestral eukaryotic microtubule networks (Fritz-Laylin et al., 2010; Wickstead et al., 2010).

Taken together, this work shows that in *Naegleria—*an internally controlled system for studying microtubule network evolution—mitotic tubulins have diversified to a remarkable extent, concurrent with the use of a limited number of regulators. This finding leads us to a model in which microtubule regulators guide the evolutionary diversification of microtubule networks for two reasons. First, and more obviously, the use of distinct microtubule regulators endows individual networks with distinct microtubule organization and dynamics. Second, cells generally use a common pool of tubulin subunits for multiple networks. Interactions with regulators in any network enforce conservation of the tubulin sequence and therefore limit the roles that tubulin specialization can play in microtubule network diversification. The tight conservation of α- and β-tubulins across phyla would therefore reflect multiple layers of constraint imposed by the large number of deeply conserved families of microtubule regulators. *Naegleria*’s unusual use of different tubulin pools for different microtubule networks provides an obvious case in which tubulin diversification can contribute to microtubule network specialization, as each set of tubulins only binds regulators used in a single network. Using the information gleaned from studying *Naegleria,* we also find similar patterns of tubulin sequence evolution in other lineages, hinting that these evolutionary mechanisms may be widespread. This model is a departure from textbook depictions of microtubule regulators as add-ons that modify fixed microtubule behaviors. Instead, the polymer and its regulators evolve together in networks, so that the nature of tubulin itself is inextricably linked to its binding partners.

## ACKNOWLEDGEMENTS

The authors wish to thank Iain Patten for valuable advice on writing and framing, as well as Pat Wadsworth, Madelaine Bartlett, and Sam Lord for insightful suggestions on the manuscript. The authors also thank members of the Fritz-Laylin lab for useful discussions. This work was supported by the National Institute of General Medical Sciences of the National Institutes of Health under award numbers F32GM148023 to A.S.K., K99GM147656 to K.B.V., and R35GM143039 to L.K.F.-L.; the Pew Charitable Trust Biomedical Scholar award to L.K.F.-L.; a Smith Family Foundation Excellence in Biomedical Research award to L.K.F.-L.; and an NSF CAREER Award number 2041395 to D.S. L.K.F.-L is a fellow of the Canadian Institute for Advanced Research, Fungal Kingdom: Threats and Opportunities program.

## AUTHOR CONTRIBUTIONS

Conceptualization: A.S.K. and L.K.F.-L.; Data curation: A.S.K.; Formal analysis: A.S.K.; Funding acquisition: A.S.K. and L.K.F.-L.; Investigation: all authors; Resources: R.R.; Supervision: L.K.F.-L.; Visualization: A.S.K., K.B.V., L.K.F.-L.; Writing – original draft: A.S.K. and L.K.F.-L. Writing – review and editing: all authors.

## DATA AVAILABILITY

Raw FASTQ files associated with this work have been deposited to the NCBI Sequence Read Archive (SRA) associated with BioProject PRJNA1061649, and will be publicly available upon publication.

## METHODS

### Cell culture

*Naegleria gruberi* strain NEG (ATCC 30223) were grown with *Aerobacter aerogenes* (both gifts from Chandler Fulton) according to standard procedures (Fulton, 1970). Briefly, for routine culturing, a spot of *Naegleria* cysts were inoculated onto one side of lawns of *Aerobacter aerogenes* that had been grown overnight at 28 °C on NM plates (2 g/L Bacto peptone (Difco), 2 g/L glucose, 1.5 g/L K_2_HPO_4_, 1 g/L KH_2_PO_4_, 2% (w/v) agar). Plates were sealed with parafilm and incubated at room temperature or 28 °C for 3-5 days, during which time cells spread across the plate, forming a prominent “edge” of rapidly dividing amoebae that eventually encysted. Cells were inoculated from the same cyst plate for up to 4 months. *Aerobacter* stock plates were streaked out from glycerol stocks stored at −80 °C, maintained at 4 °C for no more than 2 weeks, and used to inoculate stationary phase liquid cultures of *Aerobacter* that were kept at room temperature and used for up to 4 days after inoculation.

*N. gruberi* strain NEG-M (ATCC 30224) was grown axenically as previously described (Velle et al., 2022). Briefly cells were grown to a density of ∼ 1 x 10^6^ cells/ml in M7 media (0.362 g/L KH2PO4, 0.5 g/L Na2HPO4, 5.4 g/L glucose, 5 g/L yeast extract (Difco), 45 mg/L L-methionine, 10% fetal bovine serum) at 28 °C in stationary tissue culture flasks. Cells were passaged every 2-3 days by ∼20-fold dilution for no more than 30 passages before refreshing from liquid nitrogen stocks.

### Differentiation

The differentiation RNA samples used for RNA-Seq were the same ones as described in a previous publication (Fritz-Laylin & Cande, 2010). Briefly, 0.75 - 1.5 x 10^5^ amoebae were mixed with 150 µL stationary phase Aerobacter and incubated on PM plates overnight at 28 °C. Differentiation was induced by flooding the plates with ice-cold 2 mM Tris pH 7.2, scraping off amoebae, and removing bacteria through three rounds of centrifugation-based washing with ice-cold Tris buffer. Amoebae were resuspended in prewarmed Tris buffer and then incubated in an Erlenmeyer flask at 30 °C in a linear shaking water bath at 110 - 125 rpm. Samples were taken at intervals and fixed in Lugol’s iodine solution (Sigma) and the proportion of flagellates and amoebae in randomly selected fields of view were manually counted.

### Mitotic synchrony

Mitotic synchrony of suspension cultures of NEG and the food source bacterium *Aerobacter aerogenes* was conducted largely according to (Fulton & Guerrini, 1969). Briefly, lawns of *Aerobacter* were grown overnight on PM plates (4 g/L Bacto peptone (Difco), 2 g/L glucose, 1.5 g/L K_2_HPO_4_, 1 g/L KH_2_PO_4_, 2% (w/v) agar) and harvested by scraping into 10 mL of Tris (2 mM Tris pH 7.2) and centrifuging at 3,200 x g for 5-10 minutes. Bacteria were then thoroughly resuspended by vortexing in 20 mL per plate of Tris+Mg (2 mM Tris pH 7.2, 10 mM MgSO_4_).

Amoebae from the front of a growing edge plate were harvested into 1 mL of Tris using an inoculating loop and pelleted by centrifuging at 1,000 x g for 1 minute. Amoebae were resuspended in Tris and counted using a Moxi Z series impedance-based cell counter (Orflo Technologies). 3 x 10^3^ amoebae/mL were inoculated into the Tris+Mg+*Aerobacter* suspension, split into 10 mL aliquots in 125 mL Erlenmeyer flasks, and grown at 30 °C in a linear-shaking water bath at 110-125 rpm for 16-18 hours.

Amoebae from these overnight cultures were inoculated into freshly prepared Tris+Mg+*Aerobacter* suspension at a concentration of 0.5 - 1 x 10^5^ amoebae/mL in Erlenmeyer flasks at a ratio of 10 mL culture : 125 mL of nominal flask volume to ensure adequate aeration. Cultures were shaken at 110-125 rpm at 30°C until amoeba density reached 2 x 10^5^ cells/mL, at which time flasks were heat shocked by transfer to a second shaking water bath at 40.1 °C, as measured with a calibrated thermometer (Thermco Products Cat# ACC713BLS). After 100 minutes, flasks were returned to 30 °C. This time was recorded as 0 minutes post induction.

### Primary antibodies

Custom rabbit polyclonal antibodies were raised against *Naegleria* mitotic α-tubulin (NCBI: XP_002675064.1) using amino acids 26 - 47 as a peptide antigen (AEHAINQDGTRNIDSTNGNSDC), and antisera was affinity purified. Antibody generation and purification was performed by Pacific Immunology (Ramona, CA). This antibody was used for Westerns at ∼3 µg/mL, and for immunofluorescence at ∼6 µg/mL. The mouse monoclonal anti-α-tubulin antibody DM1A was used at ∼1 µg/mL for Westerns and immunofluorescence. Hybridoma culture supernatant containing a mouse monoclonal anti-β-tubulin antibody AA-12.1 (Developmental Studies Hybridoma Bank), raised against whole *Naegleria* axonemes (Shea & Walsh, 1987) was used at 1:200 dilution for Westerns and at 1:2 for immunofluorescence.

### Western blots

Samples of 1 - 3 x 10^6^ amoebae were collected per timepoint, washed to remove bacteria, and resuspended in 2mM Tris pH 7.2, 10 mM EDTA, 1X HALT protease inhibitor (Thermo Scientific, cat# 78430), and 1 mM PMSF. After a second wash, samples were lysed in 2x Laemmli’s without reducing agent (final concentration: 125 mM Tris pH 6.8, 140 mM SDS, 20% (v/v) glycerol) supplemented with 10 mM EDTA, 1X HALT protease inhibitor cocktail, and 1 mM PMSF, heated to 80°C for at least 5 minutes, and stored at −20 °C. Protein concentration of each sample was measured using the DC Protein Assay (Bio-Rad) following manufacturer instructions using BSA as a standard (Bio-Rad). Samples were diluted to 3 µg/µL in 2X Laemmli’s + 20% β-mercaptoethanol, and 15 µg of each sample was loaded into a Tris-Glycine 4-20% Stain-Free polyacrylamide gel (Bio-Rad). After electrophoresis, the stain-free reagent was activated by UV light for 5 minutes in a G:Box XX9 gel imager (Syngene). Gels were then equilibrated in Towbin’s transfer buffer without methanol (25 mM Tris, 192 mM glycine, pH 8.3) and transferred onto PVDF membranes (Amersham Hybond P Low Fluorescence) with a tank transfer system (Bio-Rad) overnight at 20 V. Transfer was confirmed by Stain-Free signal or Ponceau S staining. Blots were blocked for 30 min in TBS-T (20 mM Tris pH 7.5, 137 mM NaCl, 0.5% (v/v) Tween-20) + 5% skim milk (Difco), probed with primary antibody for 1 hr, thoroughly washed in TBS-T, and probed for 1 hr with ∼ 0.1 µg/mL horseradish peroxidase-conjugated secondary antibodies at 1:10,000 dilution (Thermo Fisher Cat# G-21234 and G-21040) diluted in TBS-T + 5% milk. After thorough washing in TBS-T, blots were imaged using freshly prepared enhanced chemiluminescent reagent (Amersham ECL Prime Cat# RPN2232) on a G:Box XX9 gel imager using 2×2 binning and varying exposures. Total protein signal (either Ponceau S or stain-free fluorescence) was quantified in ImageJ (Schindelin et al., 2012), and tubulin band intensities were quantified in Image Lab (Bio-Rad) using rolling ball background subtraction. Band intensities were normalized first to the total protein signal in each lane, and then to the intensity of the initial timepoint.

### Immunofluorescence and determination of mitotic index

Mitotic index was determined by counting spindles via immunofluorescence. Specifically, ∼ 2 x 10^5^ cells were collected, washed by centrifugation, and fixed in paraformaldehyde (25 mM sodium phosphate buffer pH 7.1, 62.5 mM sucrose, 1.8% paraformaldehyde) for 15-30 minutes at room temperature while adhering to glass coated with 0.1% polyethyleneimine. Cells were washed twice with PEM buffer (100 mM PIPES pH 6.9, 1 mM EGTA, 0.1 mM MgSO_4_), permeabilized for 10 minutes in PEM + 0.1% NP-40 alternative + 6.6 nM Alexa Fluor 488 phalloidin, and then blocked for 1 hr at room temperature in KPL Detector Block with 1% solids (SeraCare). Cells were stained overnight at 4°C with anti-α-tubulin antibody DM1A in Detector Block supplemented with 6.6 nM AF488-phalloidin. After washing 3x with Detector Block, samples were stained with anti-mouse secondary antibodies (Invitrogen) in Detector Block + 66 nM AF488-phalloidin + 1 µg/mL DAPI for 1 hr, washed 3x with Detector Block and 3x with PEM + 0.01% (v/v) NP-40 alternative, mounted in Prolong Gold + DAPI, and cured overnight at room temperature.

To visualize tubulin in amoebae and flagellates (as in Figure 2B), two samples of ∼10^5^ cells each of *N. gruberi* strain NEG-M were collected from a flask and transferred to 1.5 mL of either fresh media or 2 mM Tris (to induce differentiation) in a 6 well plate. After 1 hour of incubation, flagellates were readily observed in the Tris condition, and cells were collected and centrifuged at 1,500 x g for 90 s. Pellets were resuspended in 400 µL 2 mM Tris, and an equal volume of 2X paraformaldehyde fixative (see 1X recipe above) was immediately added. Cells were incubated for 15 min, then 200 µL of cells were added to glass coverslips coated in ∼0.1% polyethyleneimine and incubated an additional 15 min. Coverslips were rinsed twice in PEM, permeabilized for 10 min in PEM + 0.1% NP-40 alternative + 6.6 nM AF488-phalloidin, and then blocked for 1 h at room temperature in PEMBALG (PEM buffer + 1% bovine serum albumin, 100 mM lysine, 0.5% cold water fish gelatin, 0.1% sodium azide). Cells were then treated with primary antibodies (custom rabbit mitotic α-tubulin and/or mouse monoclonal anti-β-tubulin, see above) diluted in PEMBALG for 1 h, rinsed thrice in PEMBALG, and incubated for 1 h with Alexa Fluor 555 anti-mouse and/or Alexa Fluor 647 anti-rabbit secondaries (Invitrogen) diluted 1:500 in PEMBALG + 66 nM AF488-phalloidin. Cells were rinsed thrice with PEMBALG, then thrice with PEM, mounted in Prolong Gold + DAPI and cured overnight at room temperature.

Cells were imaged on a Nikon Ti2 microscope equipped with a Crest X-Light V2 L-FOV spinning disk (50 µm pinhole), a Prime95B sCMOS camera (Photometrics), a Plan Apo λD 40x 0.95 NA air objective, and a Plan Apo λ 100x 1.45 NA oil objective. The 40x objective was used with epifluorescence for calculation of the mitotic index in Figure 2D, while the 100x objective was used with spinning-disk confocal for the images in Figure 2B. Illumination was provided for epifluorescence by a Sola light source (Lumencor) and for spinning-disk confocal by a Celesta Light Engine solid-state laser launch (Lumencor) equipped with 1W lasers at 405 nm, 488 nm, 545 nm, and 637 nm. The following filter sets (all from Nikon) were used for epifluorescence imaging: ET-DAPI (excitation 378/52 nm, dichroic 409 nm, emission 447/60 nm), ET-GFP (emission 466/40 nm, dichroic 495 nm, emission 525/50 nm), and ET-DsRed (excitation 554/23 nm, dichroic 573 nm, emission 609/54 nm). The following filters were used for confocal imaging (all from Chroma unless otherwise mentioned): For DAPI, 425lpxr dichroic and ET460/50m emission; for AF488, ZT488rdc-UF1 dichroic and ET535/70m emission; for AF555, ZT561rdc-UF1 dichroic and ET610/75m emission; for AF647, FF421/491/567/659/776-Di01-25×36 dichroic (Semrock) and FF02-685/40-25 emission (Semrock). The microscope was controlled with NIS Elements.

To prevent selection bias in measurement of the mitotic index, fields of view were selected using brightfield only and therefore blinded to mitotic stage. Images were blinded and cells and spindles were counted manually in Fiji, facilitated by custom scripts. The mitotic index was computed as the percentage of cells with spindles, with at least 96 cells (and typically over 200 cells) counted per timepoint.

### RNA isolation and library preparation

RNA isolation from the differentiation experiment was previously described (Fritz-Laylin & Cande, 2010). Briefly, 1 x 10^7^ cells per timepoint were lysed in TRIzol reagent (Invitrogen), and RNA purified using RNeasy (Qiagen), treated with Turbo DNAse (Ambion), and purified a second time with RNeasy, following manufacturer instructions. RNA purity was assessed by gel electrophoresis and spectrophotometry. To isolate RNA from the mitotic synchrony samples, 4 - 8 x 10^5^ amoebae were collected per timepoint and washed free of bacteria by two rounds of vortexing and centrifugation. The amoebae were then lysed in 1 mL of TRIzol reagent and maintained at −20°C until further processing. Samples were thawed at room temperature and vortexed, and purified according to the TRIzol manufacturer’s instructions until after the chloroform extraction step, when RNA was further purified using the RNA Clean & Concentrator kit with in-column DNase-I treatment (Zymo Research Corp), following manufacturer instructions. RNA was stored at −80°C till further processing. RNA concentration was measured using the Qubit Broad Range (BR) assay kit (Life Technologies Corp) and RNA quality analyzed using Agilent 2100 Bioanalyzer RNA 6000 Nano Reagent using the Eukaryote Total RNA Nano Series II assay (Agilent Technologies Inc).

Total RNA (∼ 500 ng) was enriched for poly(A) mRNA transcripts using the NEBNext Poly(A) mRNA Magnetic Isolation Module, and libraries prepared using the NEBNext UltraII Directional RNA Library Prep Kit for Illumina (New England Biolabs) following manufacturer instructions. Library abundance was measured using the Qubit dsDNA HS (High Sensitivity) Assay Kit (Life Technologies Corp), and library quality was analyzed on an Agilent 2100 Bioanalyzer (Agilent Technologies Inc). The libraries were pooled and sequenced on Illumina NextSeq 500 platform using NextSeq 500/550 High Output v2.5 kit (150 cycles) with 76 bp paired-end sequencing chemistry. The RNA isolation and quality assessment, RNAseq library preparation and Next-Generation Sequencing (NGS) was performed at the Genomics Resource Laboratory (RRID:SCR_017907), Institute for Applied Life Sciences, University of Massachusetts Amherst, MA.

### Identification of microtubule network components in *Naegleria*

Pfam domains associated with microtubule- or tubulin-binding were identified by combining: (1) an exhaustive search of GO terms associated with microtubule binding or tubulin binding using QuickGO on 2022-10-22 (Binns et al., 2009) which were mapped to Pfam accession numbers; (2) manual searches on the InterPro website for Pfam domains that included “microtubule” or “tubulin” in their description followed by manual curation to identify domains that are directly involved with microtubules; and (3) a HHMER-based search of the translated peptide sequences of previously identified microtubule associated genes (Fritz-Laylin et al., 2010). Here, collected sequences were searched for Pfam domains in the Pfam-A v35.0 database (Mistry et al., 2021) using the hmmscan function from HMMER v3.3.2 with gathering thresholds applied. Matching domains were manually curated to exclude generic domains or other domains not directly related to microtubule- or tubulin-binding. The Pfam domains identified through these three approaches were combined to create a list of microtubule-associated Pfam domains (**Supplemental Data 4**).

To identify *Naegleria* microtubule network components, Pfam domains from the curated list described above were identified in the *Naegleria* genome using hmmsearch from HMMER 3.3.2 using gathering thresholds. The resulting hits were combined with previously identified microtubule associated genes (Fritz-Laylin et al., 2010) and additional sequences identified in the taxa-wide analysis described below, culminating in a final list of 163 *Naegleria* microtubule network components (annotated in **Supplemental Data 2**).

### Identification of microtubule network components across taxa

A list of microtubule network components was generated for homolog identification across taxa. This list (**Figure 1** and **Supplemental Data 1**) includes gene families that are well-represented in the literature and/or are defined by the presence of a family-specific, “diagnostic” Pfam domain—e.g. the kinesin motor domain, PF00225. It should be made clear that this list likely excludes less studied genes, genes that lack family-specific Pfam domains, and genes whose microtubule interactions rely on low complexity protein sequences e.g. coiled-coil or intrinsically disordered proteins that are not useful for determining homology. Homologs of the genes of interest were then identified in translated predicted proteomes for diverse species (**Supplemental Data 1**), largely by matches to the diagnostic Pfam domain(s), using hmmsearch with gathering thresholds (HMMER v3.3.2) and Pfam domains defined in Pfam-A v35.0. Hits in *Arabidopsis*, *Dictyostelium, Drosophila*, and human were manually checked against the appropriate model organism databases to validate the search strategy and to ensure only one isoform per gene locus was counted. Some gene families required modifications of this standard strategy, summarized below and also enumerated in Supplemental Data X.

Families containing functionally distinct subfamilies— tubulins, γ-tubulin complex proteins (GCPs), dyneins, and kinesins—could not be disambiguated by Pfam domain identification alone. In these cases, gene tree estimation was used to delineate subfamilies. To this end, matching sequences from all species were filtered to include only one protein product per gene locus and to remove identical sequences. Sequences passing these filters were aligned using MAFFT v7.520 with the E-INS-i algorithm and the BLOSUM45 matrix (Katoh & Standley, 2013). Gene trees were estimated from these alignments with IQ-TREE v2.2.2.7 using the LG+F+R10 substitution model (Nguyen et al., 2015). Node support was tested three ways: the ultrafast bootstrap with 10,000 replicates and 2,000 maximum iterations, the SH approximate likelihood ratio test with 1,000 replicates, and the approximate Bayes test (Hoang et al., 2018). Clades representing subfamilies were then identified by manual inspection of each tree. To distinguish among the numerous kinesin subfamilies, rather than generate an alignment *de novo*, sequences matching the kinesin motor domain (Pfam domain PF00225) from *Naegleria fowleri*, related Discoban species, and *Salpingoeca rosetta* were added to an alignment from a previous phylogenetic analysis of kinesins (Wickstead et al., 2010) using MAFFT. A kinesin gene tree was then re-estimated with IQ-TREE using the same settings and tests of support described above.

10 out of 69 of gene families (**Supplemental Data 1**) could not be reliably identified by a diagnostic Pfam domain. In the case of MAP215, a TOG domain profile from the SMART database (SM01349) was used to search the target proteomes and hits were manually inspected for the characteristic arrangement of N-terminal tandem TOG domains. For the remaining gene families, BLASTP searches were using known homologs “baits” and the set of predicted proteomes for the organisms of interest as the search space; for these searches the following settings were used: BLOSUM45 scoring matrix, word size of 3, gap open penalty of 15, gap extend penalty of 2, E-value cutoff of 1E-3. Hits were filtered to include only reciprocal hits, i.e. those that return the the query sequence as a hit when used as a query of a separate BLAST search. The resulting hits were manually curated.

An additional graph-based filtering step was performed for the microtubule severing proteins in the spectacularly large AAA-ATPase family (katanin p60, spastin, and fidgetin). To this end, a weighted, directed graph was generated from all the BLASTP hits to a Katanin p60 query sequence (human Katanin p60, Ensembl accession: , ENSP00000356381.2) with each edge defined by and weighted by the bitscore of a BLASTP hit. The subgraph of sequences connected to the bait sequence (human Katanin p60, ENSP00000356381.2) was isolated and partitioned into communities of highly interconnected hits using the Louvain community detection algorithm as implemented in the Python networkx library (Hagberg et al., 2008). Only one of the resulting communities contained the bait sequence as well as known katanin homologs from *Drosophila* and *Arabidopsis*, and so all the sequences in that community were selected for gene tree estimation as described above.

### Transcript quantification and differential expression analysis

Illumina adapter sequences were trimmed on Illumina BaseSpace. To assess the quality of the RNA-Seq coverage, reads were aligned to the 41 Mbp reference genome `(Naegr1_scaffolds.fasta` downloaded from https://phycocosm.jgi.doe.gov/Naegr1/Naegr1.home.html) using STAR v2.7.9a with default settings (Dobin et al., 2013). An example command line call is `STAR –-runThreadN 8`

`--genomeDir <path/to/genome> --readFilesCommand zcat --readFilesIn`

`<path/to/fastq.gz> --outFileNamePrefix <output/path>`

`--outReadsUnmapped Fastx --outSAMtype BAM Unsorted`. Overall, 25.5 ± 1.1 million reads per sample (mean ± s.d.) in the differentiation experiment sample and 5.1 ± 0.4 million reads per sample in the mitosis experiment were uniquely mapped to the 15,798 genes in the 24 Mbp transcriptome, resulting in 92 ± 2 % of *Naegleria* genes in the differentiation experiment and 83 ± 2 % of *Naegleria* genes in the mitosis experiment having at least one estimated transcript count (described below). To aid in correcting mis-annotated genes, read pileups for 240 transcripts of microtubule- and mitosis-associated genes from the published transcriptome were visualized with IGV (Thorvaldsdóttir et al., 2013), and 68 gene models were manually adjusted to conform to the RNA-Seq coverage across different conditions. A GFF file containing the 68 updated models (including new accessions AKG0001 - AKG0064) and with 67 superseded models removed is included as **Supplemental Data 5**. Alignment quality was further assessed using the BAMQC and RNASeqQC tools of Qualimap 2 (Okonechnikov et al., 2016).

Transcript abundance was quantified with RSEM v1.3.3 (Li & Dewey, 2011), using STAR to align reads to the transcriptome and using standard settings for Illumina paired-end sequencing. An example command line call is `rsem-calculate-expression -p 8 –-star`

`–-star-gzipped-read-file –-no-bam-output –-strandedness reverse`

`–-paired-end <path/to/upstream/reads.fastq.gz>`

`<path/to/downstream/reads.fastq.gz> <path/to/genome/indexes>`

`<path/to/output.rsem>`. Because the *Naegleria* genome has relatively few introns, and there is less than one intron per gene on average (Fritz-Laylin et al., 2010), only one transcript per gene locus was quantified and alternative splicing was not considered. The STAR aligner reports multiple possible alignments (with an associated score) for non-uniquely mapped reads (Dobin et al., 2013). These non-uniquely mapped reads made up only 7.8 ± 0.9 % of all aligned reads in each sample (mean ± s.d.). RSEM uses a generative model of RNA-Seq reads and an expectation maximization algorithm to efficiently estimate transcript counts using both uniquely and non-uniquely mapping reads (Li et al., 2010). The estimated counts generated by RSEM were fed into the DESeq2 algorithm for differential expression analysis, facilitated by the tximport function (Love et al., 2014; Soneson et al., 2016). Differential expression was tested two ways. First, differential expression was tested across the entire timecourse using a likelihood ratio test, comparing a model with time as the only factor to a constant model.

Second, differential expression was assessed post-hoc at individual timepoints using the apeglm package, which estimates adjusted log_2_ fold-changes that preserve large fold-changes while shrinking highly variable fold-changes or fold-changes from lowly expressed genes (Zhu et al., 2019). These adjusted log_2_ fold-changes were used for plotting changes in gene expression over time. Statistical significance for these fold-changes was measured using an s-value, which reflects the probability that an estimated log_2_ fold-change is the wrong sign, or is smaller than a threshold log_2_ fold-change of ±1 (Stephens, 2017).

To perform principal component analysis to visualize biological replicates, transcript counts were first normalized using the standard size factors approach implemented in DESeq2. These values were then transformed to a compressed scale using the variance stabilizing transform as implemented in the vst function of DESeq2 (Love et al., 2014). These values were finally rescaled such that the expression of each gene across all replicates and timepoints had mean 0 and variance 1. PCA was performed using the prcomp function in R.

### Clustering of expression profiles

Genes that met the following five criteria were selected for clustering, with gene sets assessed independently for each RNAseq dataset: (1) the gene was significantly differentially expressed based on the likelihood ratio test (Benjamini-Hochberg-corrected p-value < 0.05); (2) the absolute value of the adjusted log_2_ fold-change was greater than 1 for at least one timepoint; (3) the s-value associated with that log_2_ fold-change was statistically significant, i.e. less than 0.05; (4) the maximum difference in vst-normalized expression between replicates of the initial timepoint was less than a threshold value. This fourth criteria excluded genes with high expression variability across biological replicates. Due to the difference in sequencing depth between the differentiation and mitosis experiments, the absolute values of the vst-transformed transcript abundances differed, and so the threshold for “substantial” variability also differed between datasets. The threshold was chosen for each dataset to exclude a comparable number (∼100) of the most highly variable genes; (5) a fifth selection criterion was applied that was specific to the unique expression profiles in each dataset. This dataset-specific filtering was expected to exclude genes whose expression profile *a priori* were unlikely to be associated with microtubule network assembly. In the differentiation dataset, genes were included if they were *maximally* expressed *after* the initial timepoint; in the mitosis dataset, genes were included if their *minimal* expression did *not* occur at 10, 20, 30, or 40 minutes post induction of mitosis. To identify gene expression patterns among the selected genes, variance-stabilizing-transformed expression values over time were clustered using the time-series aware clustering algorithm DPGP (McDowell et al., 2018) using an α value of 0.2. The clusters returned by DPGP were further grouped using hierarchical clustering implemented in the ComplexHeatmap function in R (Gu, 2022). Clusters of interest were manually identified, with the expression of “differentiation” genes being low at 0 minutes post induction and increasing thereafter, and the expression of “mitosis” genes being low at 0 minutes post induction and maximal between 10-60 minutes post induction.

### Sequence analysis of kinesin-14 and MAP215 paralogs

The locations of domains within homologs were identified using hmmscan of the Pfam-A v35.0 database with gathering thresholds applied. TOG domains were identified with hmmscan of the TOG domain model from the SMART database (SM01349) against homologs. In all cases the predicted envelope coordinates of the domain hit were used to place the domain within the sequence. Coiled-coil domains were predicted using the Waggawagga server, and the consensus prediction from multiple algorithms was assessed manually (Simm et al., 2015). To calculate the isoelectric point of kinesin-14 homologs, each sequence was split into subsequences of 10 amino acids, and the isoelectric point of each subsequence was predicted using the `IsoelectricPoint` module in Biopython v1.78 (Cock et al., 2009).

### Tubulin sequence analysis

To identify divergent residues in mitotic tubulin, tubulin sequences from related Heteroloboseans (**Supplemental Data 1, Supplemental Figure S4**) were collected as described above as well as from a previously published tubulin gene tree (Velle et al., 2022). As previously described (Velle et al., 2022), Heterolobsean tubulin sequences fell into two clades: one containing *Naegleria gruberi* flagellate tubulins that grouped closely with tubulins from other species and another, more distant outgroup that included the *Naegleria gruberi* mitotic tubulins. Other Heterolobosean tubulins were therefore categorized as putatively flagellate or mitotic based on the clade in which they were found. For the TwinCons analysis, separate alignments of Heterolobosean α- and β-tubulins were generated with MUSCLE using the default settings on the EMBL-EBI web interface (Madeira et al., 2022). To facilitate comparisons with solved structures of pig brain microtubules, the *Sus scrofa* TUBA1B and TUBBA sequences were also included in the appropriate alignments. Per-residue conservation scores were calculated from these alignments (minus the pig tubulins) using TwinCons with the LG substitution matrix (Penev et al., 2021), comparing the putative mitotic tubulins to putative flagellate tubulins. An example command line call is `TwinCons.py -o <output/prefix> -a`

`<path/to/alignment> -lg -csv.`

Tubulin divergence in flagellated vs. non-flagellated species was calculated similarly. Specifically, tubulins from flagellate and non-flagellate species from our previous analysis were chosen; to avoid bias from overrepresentation of one species, no more than two α- and two β-tubulins were randomly selected from each species. Flagellated species in this analysis were *Batrachochytrium dendrobatidis*, *Chlamydomonas reinhardtii*, *Homo sapiens*, *Tetrahymena thermophila*, and *Trypanosoma brucei*; non-flagellated species were *Arabidopsis thaliana*, *Dictyostelium discoideum*, *Entamoeba histolytica*, *Saccharomyces cerevisiae*, and *Schizosaccharomyces pombe*. Heterolobosean species were specifically excluded from this analysis. Alignments of α- and β-tubulins were generated with MUSCLE and the TwinCons score comparing tubulins from flagellated and non-flagellated species was calculated as for the Heterolobosean flagellar/mitotic tubulins.

### Analysis of tubulin residues based on location within a microtubule

To determine how tubulin sequence divergence varies with location within a microtubule, each surface-exposed residue in α- and β-tubulin was manually categorized as facing the outside of the microtubule (lattice) or inside of the microtubule (lumen), by inspection of a molecular structure of a microtubule (PDB ID: 6O2R) in ChimeraX (Eshun-Wilson et al., 2019; Goddard et al., 2018). Surface-exposed residues were identified using the solvent-accessible surface area (SASA), which was calculated for each residue using the *measure sasa* command in ChimeraX with the default probe radius of 1.4 Å. Interfacial residues were identified as those that were solvent-accessible in the monomers but buried in the full microtubule structure. To do this, the SASA for each residue was computed for two different molecular arrangements: individual α-and β-tubulin monomers in isolation, and the tubulin heterodimer embedded in the microtubule. “Interface” residues were then defined as those whose SASA was reduced by at least 10% in the microtubule structure relative to the isolated monomers. “Lattice” residues were defined as those which were facing the outside of the microtubule and whose SASA was at least 10% of the maximum possible SASA for that amino acid (Tien et al., 2013), and “lumen” residues similarly for the inside of the microtubule. “Internal” residues were defined as those whose SASA was less than 10% of the maximum possible SASA and which were not already categorized as an interface residue.

### Analysis of tubulin:microtubule binding protein interactions

To identify tubulin residues that potentially bind to microtubule binding proteins, the Protein Data Bank was searched for structures of microtubules or tubulin dimers in contact with microtubule binding proteins and the subset of these with homologs in *Naegleria* selected for further analysis (**Supplemental Data 3**). Potentially interacting tubulin residues were identified in ChimeraX by measuring the buried surface area between the α/β-heterodimer and the microtubule binding protein using the *measure buriedarea* command. For this analysis, only protein residues were considered (as opposed to modeled waters, ions, GTP/GDP, or other non-protein components). This command reports all residues with at least 1 Å^2^ buried due to the interaction between the microtubule and the binding protein. Because some—but not all—tubulin structures adopt a particular residue numbering convention which introduces skips in the numbering of β-tubulin residues (Nogales et al., 1998), the numbering of tubulins in each structure was manually adjusted to ensure that it was consecutive and consistent across structures prior to measuring the buried area, and all interactions were then mapped to common α/β-tubulin sequences using an alignment containing all the tubulins found in the different structures (Data S1 from (Velle et al., 2022)). Tubulin residues were then defined as “interacting” with a microtubule binding protein if the buried area upon interaction was at least 20% of the solvent-accessible surface area of that residue in the same structure when the microtubule binding protein was computationally removed. In other words, a residue was considered “interacting” if at least 20% of its typically solvent-accessible surface area is buried upon association with the microtubule binding protein.

## REFERENCES

Akhmanova, A., & Steinmetz, M. O. (2015). Control of microtubule organization and dynamics: Two ends in the limelight. Nature Reviews Molecular Cell Biology, 16(12), Article 12. 10.1038/nrm4084

Alfieri, A., Gaska, I., & Forth, S. (2021). Two modes of PRC1-mediated mechanical resistance to kinesin-driven microtubule network disruption. Current Biology, 31(12), 2495–2506.e4. 10.1016/j.cub.2021.03.034

Almeida, A. C., & Maiato, H. (2018). Chromokinesins. Current Biology, 28(19), R1131–R1135. 10.1016/j.cub.2018.07.017

Atherton, J., Luo, Y., Xiang, S., Yang, C., Rai, A., Jiang, K., Stangier, M., Vemu, A., Cook, A. D., Wang, S., Roll-Mecak, A., Steinmetz, M. O., Akhmanova, A., Baldus, M., & Moores, C. A. (2019). Structural determinants of microtubule minus end preference in CAMSAP CKK domains. Nature Communications, 10(1), Article 1. 10.1038/s41467-019-13247-6

Ayaz, P., Ye, X., Huddleston, P., Brautigam, C. A., & Rice, L. M. (2012). A TOG:αβ-tubulin Complex Structure Reveals Conformation-Based Mechanisms for a Microtubule Polymerase. Science, 337(6096), 857–860. 10.1126/science.1221698

Bannigan, A., Scheible, W.-R., Lukowitz, W., Fagerstrom, C., Wadsworth, P., Somerville, C., & Baskin, T. I. (2007). A conserved role for kinesin-5 in plant mitosis. Journal of Cell Science, 120(16), 2819–2827. 10.1242/jcs.009506

Basnet, N., Nedozralova, H., Crevenna, A. H., Bodakuntla, S., Schlichthaerle, T., Taschner, M., Cardone, G., Janke, C., Jungmann, R., Magiera, M. M., Biertümpfel, C., & Mizuno, N. (2018). Direct induction of microtubule branching by microtubule nucleation factor SSNA1. Nature Cell Biology, 20(10), Article 10. 10.1038/s41556-018-0199-8

Binns, D., Dimmer, E., Huntley, R., Barrell, D., O’Donovan, C., & Apweiler, R. (2009). QuickGO: A web-based tool for Gene Ontology searching. Bioinformatics, 25(22), 3045–3046. 10.1093/bioinformatics/btp536

Bodakuntla, S., Jijumon, A. S., Villablanca, C., Gonzalez-Billault, C., & Janke, C. (2019). Microtubule-Associated Proteins: Structuring the Cytoskeleton. Trends in Cell Biology, 29(10), 804–819. 10.1016/j.tcb.2019.07.004

Bremer, N., Tria, F. D. K., Skejo, J., & Martin, W. F. (2023). The Ancestral Mitotic State: Closed Orthomitosis With Intranuclear Spindles in the Syncytial Last Eukaryotic Common Ancestor. Genome Biology and Evolution, 15(3), evad016. 10.1093/gbe/evad016

Cavalier-Smith, T. (1978). The evolutionary origin and phylogeny of microtubules, mitotic spindles and eukaryote flagella. Biosystems, 10(1–2), 93–114. 10.1016/0303-2647(78)90033-3

Chaaban, S., & Brouhard, G. J. (2017). A microtubule bestiary: Structural diversity in tubulin polymers. Molecular Biology of the Cell, 28(22), 2924–2931. 10.1091/mbc.e16-05-0271

Chang, P., Giddings, T. H., Winey, M., & Stearns, T. (2003). Ɛ-Tubulin is required for centriole duplication and microtubule organization. Nature Cell Biology, 5(1), Article 1. 10.1038/ncb900

Chung, S., Cho, J., Cheon, H., Paik, S., & Lee, J. (2002). Cloning and characterization of a divergent α-tubulin that is expressed specifically in dividing amebae of Naegleria gruberi. Gene, 293(1), 77–86. 10.1016/S0378-1119(02)00509-7

Cock, P. J. A., Antao, T., Chang, J. T., Chapman, B. A., Cox, C. J., Dalke, A., Friedberg, I., Hamelryck, T., Kauff, F., Wilczynski, B., & de Hoon, M. J. L. (2009). Biopython: Freely available Python tools for computational molecular biology and bioinformatics. Bioinformatics, 25(11), 1422–1423. 10.1093/bioinformatics/btp163

Craske, B., & Welburn, J. P. I. (2020). Leaving no-one behind: How CENP-E facilitates chromosome alignment. Essays in Biochemistry, 64(2), 313–324. 10.1042/EBC20190073

Dacheux, D., Landrein, N., Thonnus, M., Gilbert, G., Sahin, A., Wodrich, H., Robinson, D. R., & Bonhivers, M. (2012). A MAP6-Related Protein Is Present in Protozoa and Is Involved in Flagellum Motility. PLoS ONE, 7(2), e31344. 10.1371/journal.pone.0031344

Dema, A., van Haren, J., & Wittmann, T. (2022). Optogenetic EB1 inactivation shortens metaphase spindles by disrupting cortical force-producing interactions with astral microtubules. Current Biology, 32(5), 1197–1205.e4. 10.1016/j.cub.2022.01.017

Dobbelaere, J., Su, T. Y., Erdi, B., Schleiffer, A., & Dammermann, A. (2023). A phylogenetic profiling approach identifies novel ciliogenesis genes in Drosophila and C. elegans. The EMBO Journal, 42(16), e113616. 10.15252/embj.2023113616

Dobin, A., Davis, C. A., Schlesinger, F., Drenkow, J., Zaleski, C., Jha, S., Batut, P., Chaisson, M., & Gingeras, T. R. (2013). STAR: Ultrafast universal RNA-seq aligner. Bioinformatics, 29(1), 15–21. 10.1093/bioinformatics/bts635

Dutcher, S. K., Morrissette, N. S., Preble, A. M., Rackley, C., & Stanga, J. (2002). ε-Tubulin Is an Essential Component of the Centriole. Molecular Biology of the Cell, 13(11), 3859–3869. 10.1091/mbc.e02-04-0205

Eshun-Wilson, L., Zhang, R., Portran, D., Nachury, M. V., Toso, D. B., Löhr, T., Vendruscolo, M., Bonomi, M., Fraser, J. S., & Nogales, E. (2019). Effects of α-tubulin acetylation on microtubule structure and stability. Proceedings of the National Academy of Sciences, 116(21), 10366–10371. 10.1073/pnas.1900441116

Ferenz, N. P., Gable, A., & Wadsworth, P. (2010). Mitotic functions of kinesin-5. Seminars in Cell & Developmental Biology, 21(3), 255–259. 10.1016/j.semcdb.2010.01.019

Forli, S. (2014). Epothilones: From discovery to clinical trials. Current Topics in Medicinal Chemistry, 14(20), 2312–2321.

Fritz-Laylin, L. K., & Cande, W. Z. (2010). Ancestral centriole and flagella proteins identified by analysis of Naegleria differentiation. Journal of Cell Science, 123(23), 4024–4031. 10.1242/jcs.077453

Fritz-Laylin, L. K., Prochnik, S. E., Ginger, M. L., Dacks, J. B., Carpenter, M. L., Field, M. C., Kuo, A., Paredez, A., Chapman, J., Pham, J., Shu, S., Neupane, R., Cipriano, M., Mancuso, J., Tu, H., Salamov, A., Lindquist, E., Shapiro, H., Lucas, S., … Dawson, S. C. (2010). The Genome of Naegleria gruberi Illuminates Early Eukaryotic Versatility. Cell, 140(5), 631–642. 10.1016/j.cell.2010.01.032

Fulton, C. (1970). Amebo-flagellates as Research Partners: The Laboratory Biology of Naegleria and Tetramitus. Methods in Cell Biology, 4, 341–476. 10.1016/S0091-679X(08)61759-8

Fulton, C., & Dingle, A. D. (1971). Basal bodies, but not centrioles, in Naegleria. Journal of Cell Biology, 51(3), 826–836. 10.1083/jcb.51.3.826

Fulton, C., & Guerrini, A. M. (1969). Mitotic synchrony in Naegleria amebae. Experimental Cell Research, 56(2), 194–200. 10.1016/0014-4827(69)90002-0

Fulton, C., & Simpson, P. A. (1976). Selective synthesis and utilization of flagellar tubulin. The multi-tubulin hypothesis. Cell Motility, 3, 987–1005.

Gard, D. L., Becker, B. E., & Josh Romney, S. (2004). MAPping the Eukaryotic Tree of Life: Structure, Function, and Evolution of the MAP215⧸Dis1 Family of Microtubule-Associated Proteins. International Review of Cytology, 239, 179–272. 10.1016/S0074-7696(04)39004-2

Gardiner, J. (2013). The evolution and diversification of plant microtubule-associated proteins. The Plant Journal, 75(2), 219–229. 10.1111/tpj.12189

Gassmann, R. (2023). Dynein at the kinetochore. Journal of Cell Science, 136(5), jcs220269. 10.1242/jcs.220269

Goddard, T. D., Huang, C. C., Meng, E. C., Pettersen, E. F., Couch, G. S., Morris, J. H., & Ferrin, T. E. (2018). UCSF ChimeraX: Meeting modern challenges in visualization and analysis. Protein Science, 27(1), 14–25. 10.1002/pro.3235

Goodson, H. V., & Jonasson, E. M. (2018). Microtubules and Microtubule-Associated Proteins. Cold Spring Harbor Perspectives in Biology, 10(6), a022608. 10.1101/cshperspect.a022608

Gu, Z. (2022). Complex heatmap visualization. iMeta, 1(3), e43. 10.1002/imt2.43

Guo, J., Qiang, M., & Ludueña, R. F. (2011). The distribution of β-tubulin isotypes in cultured neurons from embryonic, newborn, and adult mouse brains. Brain Research, 1420, 8–18. 10.1016/j.brainres.2011.08.066

Gupta, M. L., Bode, C. J., Georg, G. I., & Himes, R. H. (2003). Understanding tubulin–Taxol interactions: Mutations that impart Taxol binding to yeast tubulin. Proceedings of the National Academy of Sciences, 100(11), 6394–6397. 10.1073/pnas.1131967100

Hagberg, A. A., Schult, D. A., & Swart, P. J. (2008). Exploring Network Structure, Dynamics, and Function using NetworkX. In G. Varoquaux, T. Vaught, & J. Millman (Eds.), Proceedings of the 7th Python in Science Conference (pp. 11–15).

Heath, I. B. (1980). Variant Mitoses in Lower Eukaryotes: Indicators of the Evolution of Mitosis? International Review of Cytology, 64, 1–80. 10.1016/S0074-7696(08)60235-1

Higashide, E., Asai, M., Ootsu, K., Tanida, S., Kozai, Y., Hasegawa, T., Kishi, T., Sugino, Y., & Yoneda, M. (1977). Ansamitocin, a group of novel maytansinoid antibiotics with antitumour properties from Nocardia. Nature, 270(5639), Article 5639. 10.1038/270721a0

Hoang, D. T., Chernomor, O., von Haeseler, A., Minh, B. Q., & Vinh, L. S. (2018). UFBoot2: Improving the Ultrafast Bootstrap Approximation. Molecular Biology and Evolution, 35(2), 518–522. 10.1093/molbev/msx281

Hou, Y., Sierra, R., Bassen, D., Banavali, N. K., Habura, A., Pawlowski, J., & Bowser, S. S. (2013). Molecular Evidence for β-tubulin Neofunctionalization in Retaria (Foraminifera and Radiolarians). Molecular Biology and Evolution, 30(11), 2487–2493. 10.1093/molbev/mst150

Ichikawa, M., Liu, D., Kastritis, P. L., Basu, K., Hsu, T. C., Yang, S., & Bui, K. H. (2017). Subnanometre-resolution structure of the doublet microtubule reveals new classes of microtubule-associated proteins. Nature Communications, 8(1), Article 1. 10.1038/ncomms15035

Janke, C., & Magiera, M. M. (2020). The tubulin code and its role in controlling microtubule properties and functions. Nature Reviews Molecular Cell Biology, 21(6), Article 6. 10.1038/s41580-020-0214-3

Kajtez, J., Solomatina, A., Novak, M., Polak, B., Vukušić, K., Rüdiger, J., Cojoc, G., Milas, A., Šumanovac Šestak, I., Risteski, P., Tavano, F., Klemm, A. H., Roscioli, E., Welburn, J., Cimini, D., Glunčić, M., Pavin, N., & Tolić, I. M. (2016). Overlap microtubules link sister k-fibres and balance the forces on bi-oriented kinetochores. Nature Communications, 7(1), Article 1. 10.1038/ncomms10298

Katoh, K., & Standley, D. M. (2013). MAFFT Multiple Sequence Alignment Software Version 7: Improvements in Performance and Usability. Molecular Biology and Evolution, 30(4), 772–780. 10.1093/molbev/mst010

Kraus, J., Alfaro-Aco, R., Gouveia, B., & Petry, S. (2023). Microtubule nucleation for spindle assembly: One molecule at a time. Trends in Biochemical Sciences, 48(9), 761–775. 10.1016/j.tibs.2023.06.004

Lee, A. H.-Y., Hurley, B., Felsensteiner, C., Yea, C., Ckurshumova, W., Bartetzko, V., Wang, P. W., Quach, V., Lewis, J. D., Liu, Y. C., Börnke, F., Angers, S., Wilde, A., Guttman, D. S., & Desveaux, D. (2012). A Bacterial Acetyltransferase Destroys Plant Microtubule Networks and Blocks Secretion. PLOS Pathogens, 8(2), e1002523. 10.1371/journal.ppat.1002523

Lewis, S. A., Gu, W., & Cowan, N. J. (1987). Free intermingling of mammalian β-tubulin isotypes among functionally distinct microtubules. Cell, 49(4), 539–548. 10.1016/0092-8674(87)90456-9

Li, B., & Dewey, C. N. (2011). RSEM: Accurate transcript quantification from RNA-Seq data with or without a reference genome. BMC Bioinformatics, 12(1), 323. 10.1186/1471-2105-12-323

Li, B., Ruotti, V., Stewart, R. M., Thomson, J. A., & Dewey, C. N. (2010). RNA-Seq gene expression estimation with read mapping uncertainty. Bioinformatics, 26(4), 493–500. 10.1093/bioinformatics/btp692

Lopata, M. A., & Cleveland, D. W. (1987). In vivo microtubules are copolymers of available beta-tubulin isotypes: Localization of each of six vertebrate beta-tubulin isotypes using polyclonal antibodies elicited by synthetic peptide antigens. Journal of Cell Biology, 105(4), 1707–1720. 10.1083/jcb.105.4.1707

Love, M. I., Huber, W., & Anders, S. (2014). Moderated estimation of fold change and dispersion for RNA-seq data with DESeq2. Genome Biology, 15(12), 550. 10.1186/s13059-014-0550-8

Lüdecke, A., Seidel, A.-M., Braun, M., Lansky, Z., & Diez, S. (2018). Diffusive tail anchorage determines velocity and force produced by kinesin-14 between crosslinked microtubules. Nature Communications, 9(1), Article 1. 10.1038/s41467-018-04656-0

Madeira, F., Pearce, M., Tivey, A. R. N., Basutkar, P., Lee, J., Edbali, O., Madhusoodanan, N., Kolesnikov, A., & Lopez, R. (2022). Search and sequence analysis tools services from EMBL-EBI in 2022. Nucleic Acids Research, 50(W1), W276–W279. 10.1093/nar/gkac240

McDowell, I. C., Manandhar, D., Vockley, C. M., Schmid, A. K., Reddy, T. E., & Engelhardt, B. E. (2018). Clustering gene expression time series data using an infinite Gaussian process mixture model. PLOS Computational Biology, 14(1), e1005896. 10.1371/journal.pcbi.1005896

McNally, F. J., & Roll-Mecak, A. (2018). Microtubule-severing enzymes: From cellular functions to molecular mechanism. Journal of Cell Biology, 217(12), 4057–4069. 10.1083/jcb.201612104

Mistry, J., Chuguransky, S., Williams, L., Qureshi, M., Salazar, G. A., Sonnhammer, E. L. L., Tosatto, S. C. E., Paladin, L., Raj, S., Richardson, L. J., Finn, R. D., & Bateman, A. (2021). Pfam: The protein families database in 2021. Nucleic Acids Research, 49(D1), D412–D419. 10.1093/nar/gkaa913

Mitchell, D. R. (2017). Evolution of Cilia. Cold Spring Harbor Perspectives in Biology, 9(1), a028290. 10.1101/cshperspect.a028290

Mountain, V., Simerly, C., Howard, L., Ando, A., Schatten, G., & Compton, D. A. (1999). The Kinesin-Related Protein, Hset, Opposes the Activity of Eg5 and Cross-Links Microtubules in the Mammalian Mitotic Spindle. Journal of Cell Biology, 147(2), 351–366. 10.1083/jcb.147.2.351

Nguyen, L.-T., Schmidt, H. A., von Haeseler, A., & Minh, B. Q. (2015). IQ-TREE: A Fast and Effective Stochastic Algorithm for Estimating Maximum-Likelihood Phylogenies. Molecular Biology and Evolution, 32(1), 268–274. 10.1093/molbev/msu300

Nielsen, M. G., Gadagkar, S. R., & Gutzwiller, L. (2010). Tubulin evolution in insects: Gene duplication and subfunctionalization provide specialized isoforms in a functionally constrained gene family. BMC Evolutionary Biology, 10(1), 113. 10.1186/1471-2148-10-113

Nielsen, M. G., Turner, F. R., Hutchens, J. A., & Raff, E. C. (2001). Axoneme-specific β-tubulin specialization: A conserved C-terminal motif specifies the central pair. Current Biology, 11(7), 529–533. 10.1016/S0960-9822(01)00150-6

Nogales, E., Wolf, S. G., & Downing, K. H. (1998). Structure of the αβ tubulin dimer by electron crystallography. Nature, 391(6663), Article 6663. 10.1038/34465

Okonechnikov, K., Conesa, A., & García-Alcalde, F. (2016). Qualimap 2: Advanced multi-sample quality control for high-throughput sequencing data. Bioinformatics, 32(2), 292–294. 10.1093/bioinformatics/btv566

Owa, M., Uchihashi, T., Yanagisawa, H., Yamano, T., Iguchi, H., Fukuzawa, H., Wakabayashi, K., Ando, T., & Kikkawa, M. (2019). Inner lumen proteins stabilize doublet microtubules in cilia and flagella. Nature Communications, 10(1), Article 1. 10.1038/s41467-019-09051-x

Penev, P. I., Alvarez-Carreño, C., Smith, E., Petrov, A. S., & Williams, L. D. (2021). TwinCons: Conservation score for uncovering deep sequence similarity and divergence. PLOS Computational Biology, 17(10), e1009541. 10.1371/journal.pcbi.1009541

Petry, S., Groen, A. C., Ishihara, K., Mitchison, T. J., & Vale, R. D. (2013). Branching Microtubule Nucleation in Xenopus Egg Extracts Mediated by Augmin and TPX2. Cell, 152(4), 768–777. 10.1016/j.cell.2012.12.044

Pickett-Heaps, J. (1974). The evolution of mitosis and the eukaryotic condition. Biosystems, 6(1), 37–48. 10.1016/0303-2647(74)90009-4

Pickett-Heaps, J. D. (1975). ASPECTS OF SPINDLE EVOLUTION*. Annals of the New York Academy of Sciences, 253(1), 352–361. 10.1111/j.1749-6632.1975.tb19213.x

Pidoux, A. L., LeDizet, M., & Cande, W. Z. (1996). Fission yeast pkl1 is a kinesin-related protein involved in mitotic spindle function. Molecular Biology of the Cell, 7(10), 1639–1655. 10.1091/mbc.7.10.1639

Pucciarelli, S., Ballarini, P., Sparvoli, D., Barchetta, S., Yu, T., Iii, H. W. D., & Miceli, C. (2012). Distinct Functional Roles of β-Tubulin Isotypes in Microtubule Arrays of Tetrahymena thermophila, a Model Single-Celled Organism. PLOS ONE, 7(6), e39694. 10.1371/journal.pone.0039694

Roll-Mecak, A. (2020). The Tubulin Code in Microtubule Dynamics and Information Encoding. Developmental Cell, 54(1), 7–20. 10.1016/j.devcel.2020.06.008

Ruiz, F., Krzywicka, A., Klotz, C., Keller, A.-M., Cohen, J., Koll, F., Balavoine, G., & Beisson, J. (2000). The SM19 gene, required for duplication of basal bodies in Paramecium, encodes a novel tubulin, η-tubulin. Current Biology, 10(22), 1451–1454. 10.1016/S0960-9822(00)00804-6

Schindelin, J., Arganda-Carreras, I., Frise, E., Kaynig, V., Longair, M., Pietzsch, T., Preibisch, S., Rueden, C., Saalfeld, S., Schmid, B., Tinevez, J.-Y., White, D. J., Hartenstein, V., Eliceiri, K., Tomancak, P., & Cardona, A. (2012). Fiji: An open-source platform for biological-image analysis. Nature Methods, 9(7), Article 7. 10.1038/nmeth.2019

Shea, D. K., & Walsh, C. J. (1987). mRNAs for alpha- and beta-tubulin and flagellar calmodulin are among those coordinately regulated when Naegleria gruberi amebae differentiate into flagellates. Journal of Cell Biology, 105(3), 1303–1309. 10.1083/jcb.105.3.1303

Simm, D., Hatje, K., & Kollmar, M. (2015). Waggawagga: Comparative visualization of coiled-coil predictions and detection of stable single α-helices (SAH domains). Bioinformatics, 31(5), 767–769. 10.1093/bioinformatics/btu700

Soneson, C., Love, M. I., & Robinson, M. D. (2016). Differential analyses for RNA-seq: Transcript-level estimates improve gene-level inferences (4:1521). F1000Research. 10.12688/f1000research.7563.2

Stephens, M. (2017). False discovery rates: A new deal. Biostatistics, 18(2), 275–294. 10.1093/biostatistics/kxw041

Strassert, J. F. H., Irisarri, I., Williams, T. A., & Burki, F. (2021). A molecular timescale for eukaryote evolution with implications for the origin of red algal-derived plastids. Nature Communications, 12(1), Article 1. 10.1038/s41467-021-22044-z

Thorvaldsdóttir, H., Robinson, J. T., & Mesirov, J. P. (2013). Integrative Genomics Viewer (IGV): High-performance genomics data visualization and exploration. Briefings in Bioinformatics, 14(2), 178–192. 10.1093/bib/bbs017

Tian, G., & Cowan, N. J. (2013). Chapter 11 - Tubulin-Specific Chaperones: Components of a Molecular Machine That Assembles the α/β Heterodimer. In J. J. Correia & L. Wilson (Eds.), Methods in Cell Biology (Vol. 115, pp. 155–171). Academic Press. 10.1016/B978-0-12-407757-7.00011-6

Tien, M. Z., Meyer, A. G., Sydykova, D. K., Spielman, S. J., & Wilke, C. O. (2013). Maximum Allowed Solvent Accessibilites of Residues in Proteins. PLOS ONE, 8(11), e80635. 10.1371/journal.pone.0080635

Velle, K. B., Kennard, A. S., Trupinić, M., Ivec, A., Swafford, A. J. M., Nolton, E., Rice, L. M., Tolić, I. M., Fritz-Laylin, L. K., & Wadsworth, P. (2022). Naegleria’s mitotic spindles are built from unique tubulins and highlight core spindle features. Current Biology, 32(6), 1247–1261.e6. 10.1016/j.cub.2022.01.034

Walsh, C. J. (2007). The role of actin, actomyosin and microtubules in defining cell shape during the differentiation of Naegleria amebae into flagellates. European Journal of Cell Biology, 86(2), 85–98. 10.1016/j.ejcb.2006.10.003

Wickstead, B., & Gull, K. (2007). Dyneins Across Eukaryotes: A Comparative Genomic Analysis. Traffic (Copenhagen, Denmark), 8(12), 1708–1721. 10.1111/j.1600-0854.2007.00646.x

Wickstead, B., Gull, K., & Richards, T. A. (2010). Patterns of kinesin evolution reveal a complex ancestral eukaryote with a multifunctional cytoskeleton. BMC Evolutionary Biology, 10(1), 110. 10.1186/1471-2148-10-110

Zhu, A., Ibrahim, J. G., & Love, M. I. (2019). Heavy-tailed prior distributions for sequence count data: Removing the noise and preserving large differences. Bioinformatics, 35(12), 2084–2092. 10.1093/bioinformatics/bty895

