## Supplemental Figures 1-4 for "An internally controlled system to study microtubule network diversification links tubulin evolution to the use of distinct microtubule regulators"

### Figure S1

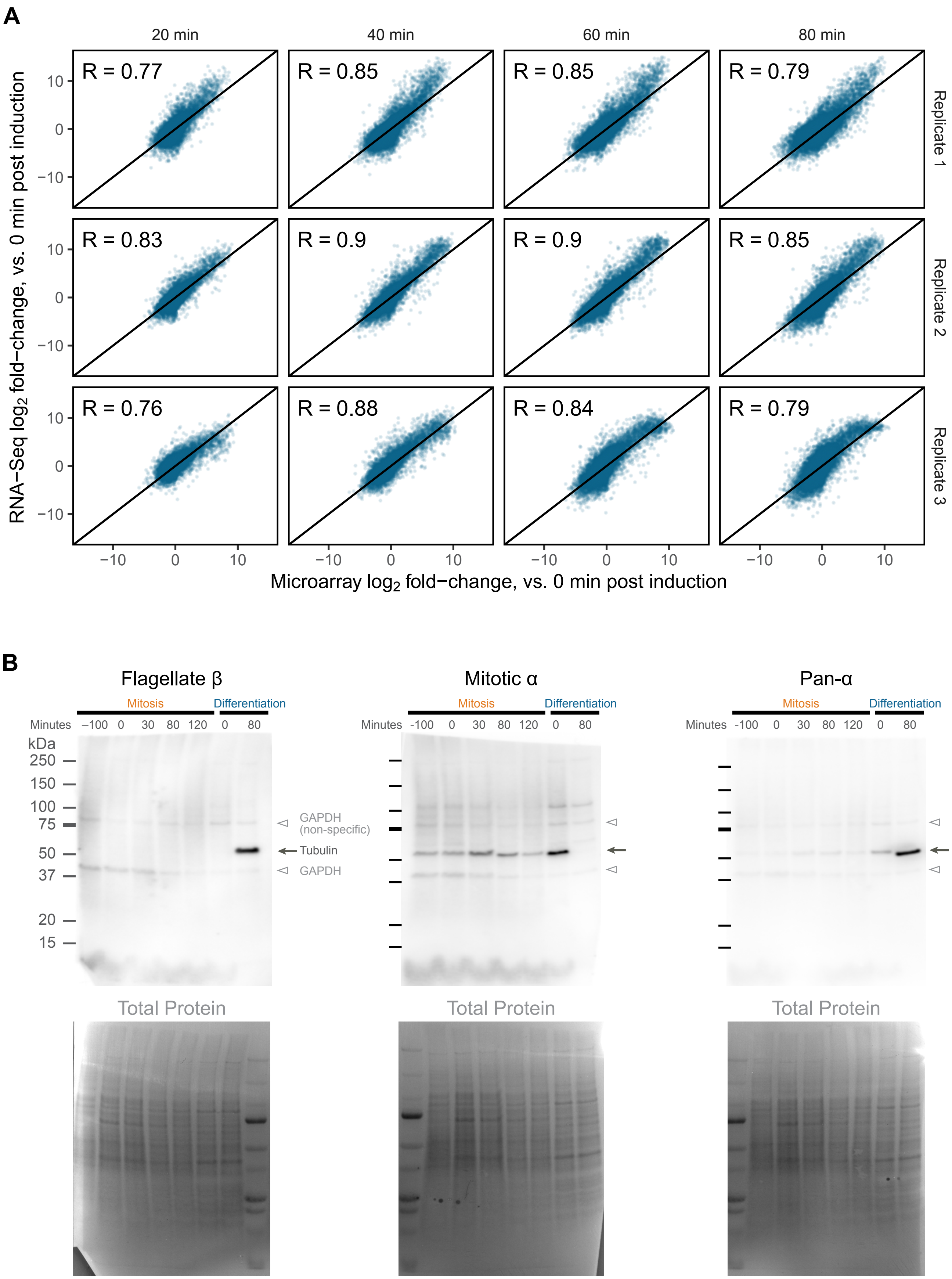
A
20 min
40 min
60 min
80 min
R = 0.77
R = 0.85
R = 0.85
R = 0.79
R = 0.83
R = 0.9
R = 0.9
R = 0.85
R = 0.76
R = 0.88
R = 0.84
R = 0.79
Replicate 1
Replicate 2
Replicate 3
RNA−Seq log2 fold−change, vs. 0 min post induction
Microarray log2 fold−change, vs. 0 min post induction
B
Flagellate β
Mitotic α
Pan-α
Mitosis
Differentiation
Minutes −100 0 30 80 120 0 80
kDa
250
150
100
75
50
37
20
15
GAPDH (non-specific)
Tubulin
GAPDH
Total Protein

**Figure S1: Related to Figure 2. (A)** Correlation of gene expression fold-changes measured by RNA-Seq and microarray of the same samples from the differentiation experiment. Microarray data was taken from Fritz-Laylin and Cande, (2010). *J Cell Sci 123*(23): 4024-4031. Microarray expression data was normalized as described in that publication, and then log2-fold changes were calculated relative to 0 minutes post induction for each timepoint. Log2 fold-changes for the RNA-Seq data were calculated directly from the sample counts normalized for library size factors in DESeq2, with no other normalization applied. Each dot represents expression of a gene in one biological replicate of one timepoint. A black line marks $y = x$ line to guide the eye. The Pearson correlation coefficient for each scatterplot is shown in the top left corner of each panel. **(B)** Full images of Western blots shown in Figure 2E. Ponceau S total protein staining is shown below each blot. The gray arrow marks the tubulin band at ~54 kDa. Arrowheads denote bands generated by a GAPDH antibody that was co-incubated with the tubulin antibodies in all three blots.

Figure S2

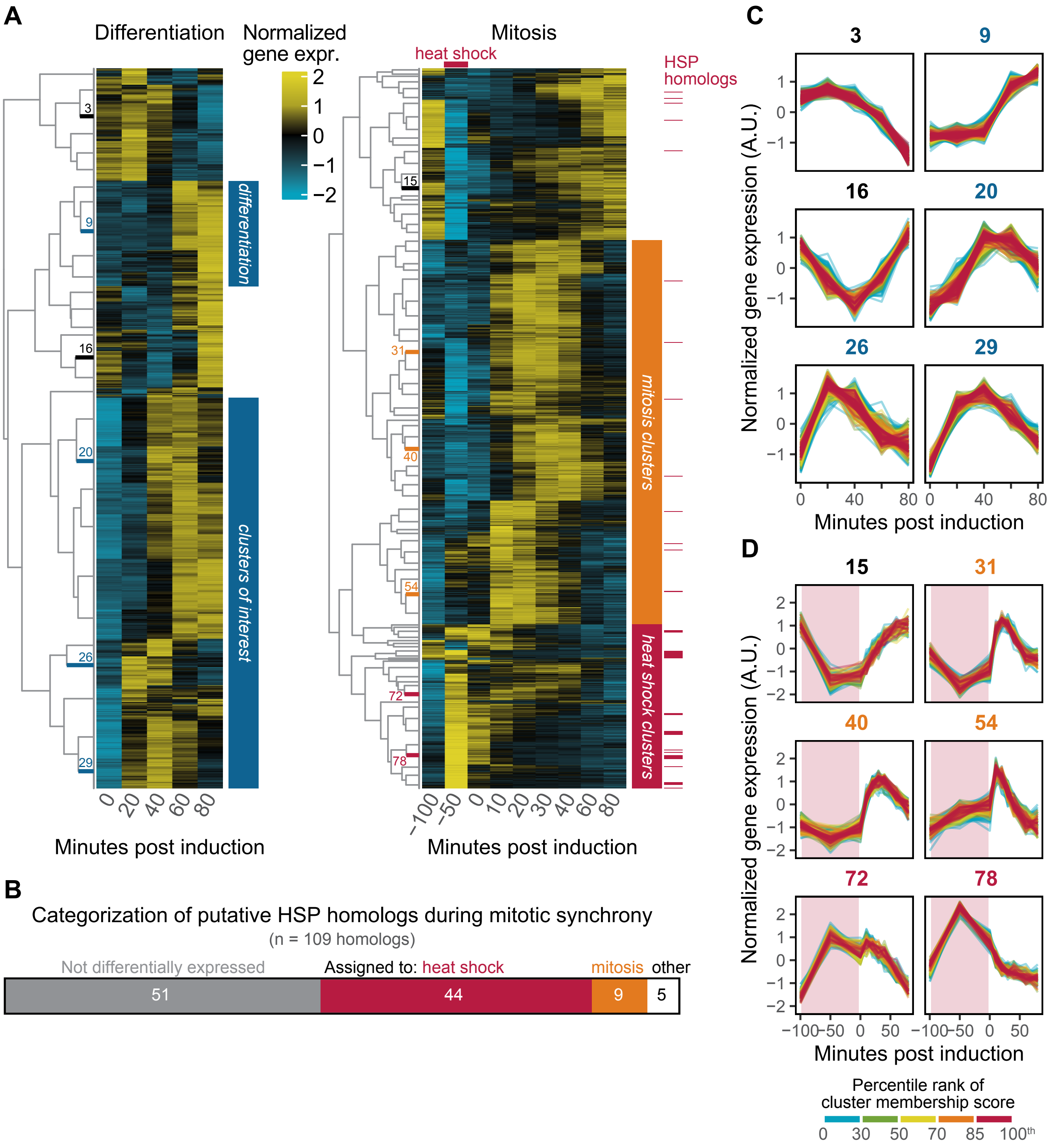
A
Differentiation
Normalized gene expr.
Mitosis
heat shock
HSP homologs
differentiation
clusters of interest
mitosis clusters
heat shock clusters
Minutes post induction
Minutes post induction
B
Categorization of putative HSP homologs during mitotic synchrony
(n = 109 homologs)
Not differentially expressed
Assigned to: heat shock
mitosis
other
51
44
9
5
C
Normalized gene expression (A.U.)
Minutes post induction
D
Normalized gene expression (A.U.)
Minutes post induction
Percentile rank of cluster membership score
0 30 50 70 85 100th

**Figure S2: Related to Figure 3. (A)** Complete heatmaps of differential gene expression in both experiments. The differentiation heatmap is reproduced from Figure 3B. The mitosis heatmap also includes two timepoints before and during the synchronizing heat shock, marked -100 and -50 minutes post induction. Homologs of heat-shock related proteins are marked on the right of the mitosis heatmap. Homologs were identified based on hits to diagnostic Pfam domains (provided in Supplemental Data 2), e.g. HSP70. The width of each mark is not to scale but is roughly proportional to the number of homologs in neighboring rows. **(B)** Categorization of the expression of heat-shock related protein homologs. Out of 109 homologs, nearly half (n = 51) were not differentially expressed in mitosis. Of the remainder, the vast majority were assigned to the heat shock clusters marked in red in (A). **(C-D)** Plots of normalized gene expression for example clusters from differentiation (C) and mitosis (D). The position of clusters in the heatmap is marked in panel (A), and the cluster numbers are color-coded based on if they are part of a marked cluster grouping (i.e. mitosis, differentiation, or heat shock). Each line represents the transcript abundance of a gene over time; to facilitate visual comparison, transcript abundance is normalized across the timecourse to mean 0 and standard deviation 1. Trajectories are colored by the percentile rank of their “membership score” within the cluster. The membership score is the posterior probability of cluster membership generated by the DPGP clustering algorithm. The red bands in (D) indicate the timing of the heat shock in the mitosis experiment.

### Figure S3

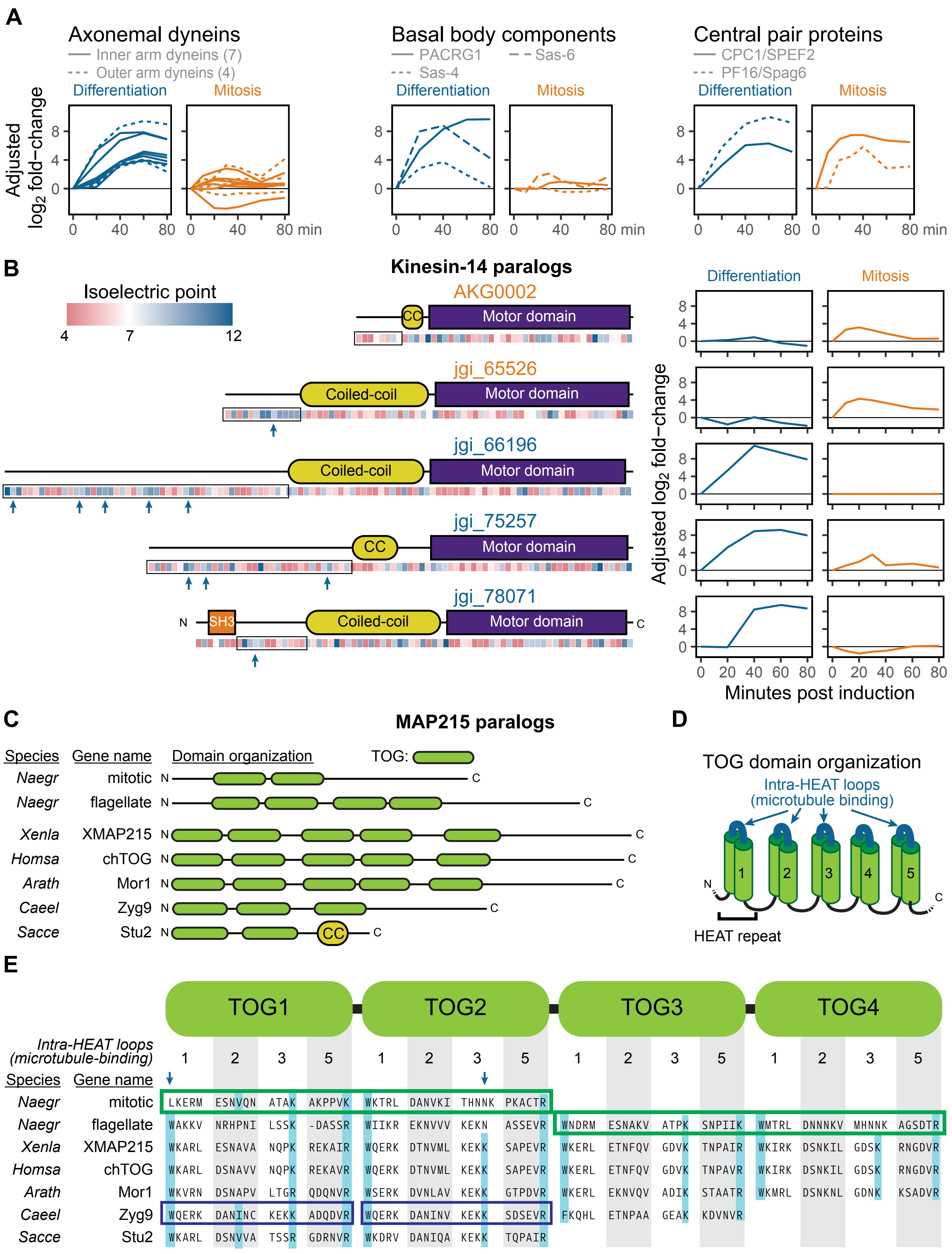

A
Axonemal dyneins
Inner arm dyneins (7)
Outer arm dyneins (4)
Differentiation
Mitosis
Adjusted log2 fold−change
0 4 8
0 40 80 0 40 80 min
Basal body components
PACRG1
Sas-6
Sas-4
Differentiation
Mitosis
Central pair proteins
CPC1/SPEF2
PF16/Spag6
Differentiation
Mitosis
B
Kinesin-14 paralogs
Isoelectric point
4 7 12
AKG0002
CC
Motor domain
jgi_65526
Coiled-coil
Motor domain
jgi_66196
Coiled-coil
Motor domain
jgi_75257
CC
Motor domain
jgi_78071
N
SH3
Coiled-coil
Motor domain
C
Differentiation
Mitosis
Adjusted log2 fold−change
0 20 40 60 80 0 20 40 60 80
Minutes post induction
C
MAP215 paralogs
Species Gene name Domain organization TOG:
Naegr mitotic N C
Naegr flagellate N C
Xenla XMAP215 N C
Homsa chTOG N C
Arath Mor1 N C
Caeel Zyg9 N C
Sacce Stu2 N CC C
D
TOG domain organization
Intra-HEAT loops (microtubule binding)
N 1 2 3 4 5 C
HEAT repeat
E
TOG1 TOG2 TOG3 TOG4
Intra-HEAT loops (microtubule-binding)
1 2 3 5 1 2 3 5 1 2 3 5 1 2 3 5
Species Gene name
Naegr mitotic LKERM ESNVQN ATAK AKPPVK WKTRL DANVKI THNNK PKACTR
Naegr flagellate WAKKV NRHPNI LSSK -DASSR WIIKR EKNVVV KEKN ASSEVR WNDRM ESNAKV ATPK SNPIIK WMTRL DNNNKV MHNNK AGSDTR
Xenla XMAP215 WKARL ESNAVA NQPK REKAIR WQERK DTNVML KEKK SAPEVR WKERL ETNFQV GDVK TNPAIR WKIRK DSNKIL GDSK RNGDVR
Homsa chTOG WKARL DSNAVV NQPK REKAVR WQERK DTNVML KEKK SAPEVR WKERL ETNFQV GDVK TNPAVR WKIRK DSNKIL GDSK RNGDVR
Arath Mor1 WKVRN DSNAPV LTGR QDQNVR WSERK DVNLAV KEKK GTPDVR WKERL EKNVQV ADIK STAATR WKMRL DSNKNL GDNK KSADVR
Caeel Zyg9 WQERK DANINC KEKK ADQDVR WQERK DANINV KEKK SDSEVR FKQHL ETNPAA GEAK KDVNVR
Sacce Stu2 WKARL DSNVVA TSSR GDRNVR WKDRV DANIQA KEKK TQPAIR

**Figure S3: Related to Figure 4. (A)** Expression of key axonemal components in differentiation and mitosis. Adjusted log2 fold-change (see Methods) over time for different categories of genes, including axonemal dyneins (*left*), proteins involved in basal body formation (*middle*), and proteins associated with the central pair (*right*). Homology for the basal body and central pair proteins was identified in Fritz-Laylin *et al.* (2010) *Cell 140*(5): 631-642. **(B)** Domain analysis of *Naegleria* kinesin-14 paralogs. Domains identified by Pfam and predicted coiled-coil regions (see Methods) are shown to scale for each homolog. The isoelectric point computed locally in 10-amino-acid windows is displayed below each domain map. Blue arrows indicate contiguous basic sequence regions of at least 20 amino acids. The gene identifiers are color-coded based on whether the paralog is categorized as differentiation- or mitosis- specific. Expression of each homolog in differentiation and mitosis is shown at right. **(C)** Domain maps of *Naegleria* MAP215 paralogs and homologs in model organisms, including *Xenopus laevis*, *Homo sapiens*, *Arabidopsis thaliana*, *Caenorhabditis elegans*, and *Saccharomyces cerevisiae*. The *Saccharomyces* homolog Stu2 also has a coiled-coil region that was not detected in other homologs. **(D)** Intra-domain schematic of TOG domains, which are formed of five tandem HEAT repeats separated by intra-HEAT loops. These loops contain highly conserved residues that are essential for binding to tubulin dimers. **(E)** Alignments of the intra-HEAT loops from the different TOG domains of the homologs shown in (C). Teal bars indicate highly conserved residues involved in tubulin binding (cf. Ayaz *et al.* (2014) *eLife 3*: e03069). Blue arrows point to examples of such positions that are not conserved in the *Naegleria* mitotic MAP215 paralog. The purple boxes illustrate a likely TOG domain duplication within *C. elegans* Zyg9, in which the TOG1 domain of Zyg9 was likely formed by duplication of the TOG2 domain. Green boxes show possible homology between TOG3-TOG4 of the *Naegleria* flagellate MAP215 paralog and TOG1-TOG2 of the mitotic paralog.

Figure S4

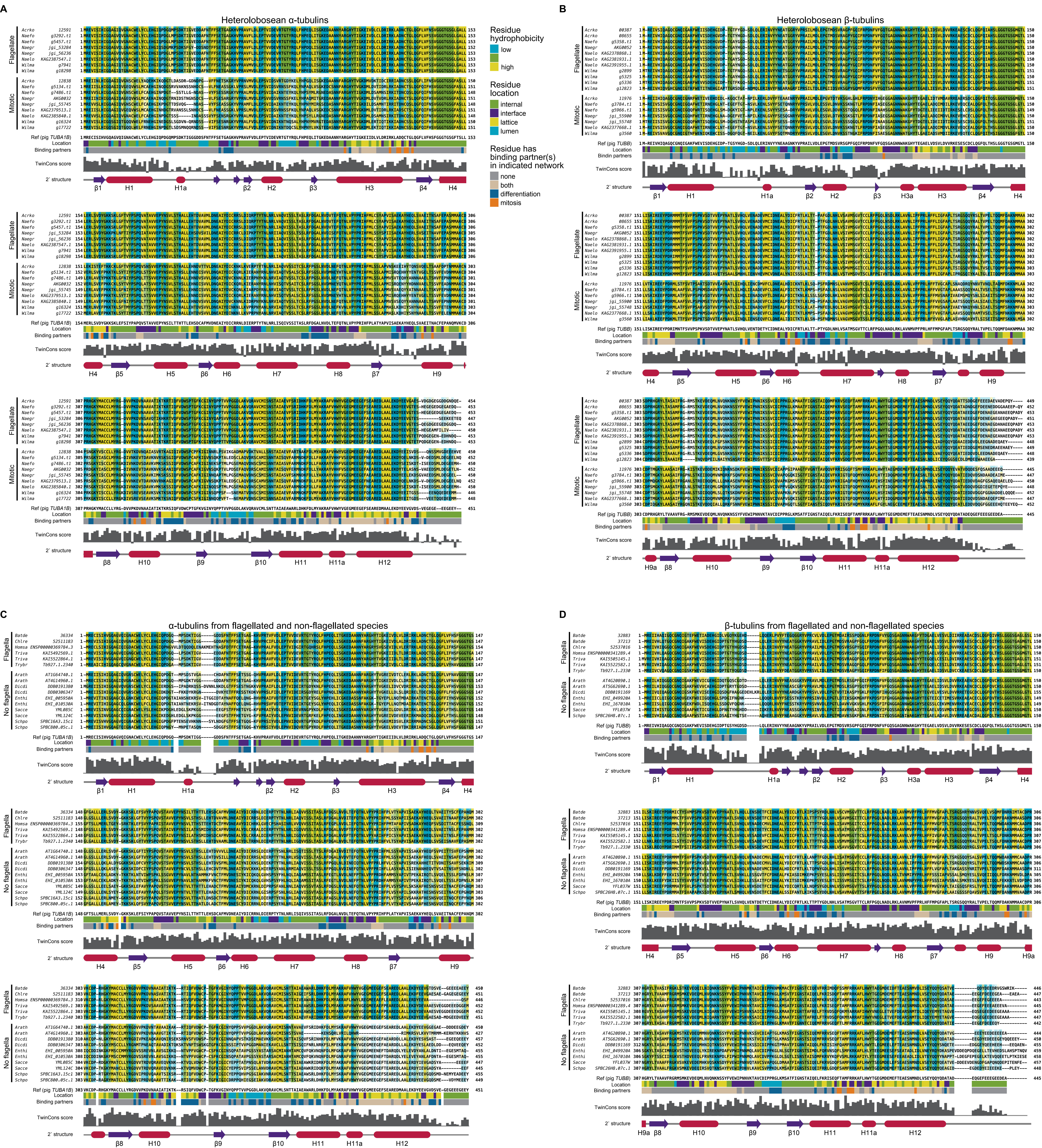
A
Heterolobosean α-tubulins
B
Heterolobosean β-tubulins
Flagellate
Mitotic
Residue hydrophobicity
low
high
Residue location
internal
interface
lattice
lumen
Residue has binding partner(s) in indicated network
none
both
differentiation
mitosis
Ref (pig TUBA1B)
Ref (pig TUBB)
Location
Binding partners
TwinCons score
2' structure
C
α-tubulins from flagellated and non-flagellated species
D
β-tubulins from flagellated and non-flagellated species
Flagella
No flagella

**Figure S4: Related to Figure 5. (A-B)** Alignments of heterolobosean α-tubulins (A) and β-tubulins (B), split into mitotic and flagellate groups. Residues are colored according to relative hydrophobicity, and shaded based on conservation *within* each group of the alignment. The pig TUBA1B and TUBB sequences are shown for reference, as well as the classification of each residue's location in the microtubule, and whether each residue is predicted to have binding partners in mitosis, differentiation, both, or neither. The TwinCons score calculated from each alignment is shown, as well as the secondary structure, extracted from the assignments in the PDB file of a pig microtubule structure (PDB: 6O2R), following standard secondary structure naming conventions for tubulin. **(C-D)** Alignments formatted as for (A) and (B), but with sequences originating from species that have flagella or do not have them. Gaps in the tracks correspond to positions in which the reference tubulin has gaps.
